# Energetic niches of syntrophic methanogens recorded by clumped isotopes

**DOI:** 10.64898/2026.11.14.766673

**Authors:** Jiawen Li, Laetitia Guibourdenche, Dari Bumgarner, Michael Galvin, Annie Kandel, Xiahong Feng, Anna Schnürer, Maria Westerholm, Anna Martini, James F. Holden, Edward D. Young, William D. Leavitt

## Abstract

Methane is a potent greenhouse gas, an important energy source and a potential biosignature in extraterrestrial environments. Stable carbon and hydrogen isotope (*δ*^13^C and *δ*D) and clumped isotopologues (Δ^13^CH_3_D and Δ^12^CH_2_D_2_) compositions of methane have been widely used as tracers for methanogenesis pathways. Furthermore, isotopic signatures of methane typically shift closer to equilibrium as the Gibbs energy (ΔG_r_) of methanogenesis moves closer to the energy limit of life. However, the variations of clumped methane isotope signatures in response to the changes in ΔG_r_ lack careful examination. In this study, we measure the isotope signatures of methane produced by monoculture methanogens with formate or H_2_/CO_2_, and by syntrophic cocultures of hydrogenotrophic methanogens paired with acetate or propionate oxidizers. Methanogenesis by cocultures has lower Gibbs energy yield than monocultures, yet both Δ^13^CH_3_D and Δ^12^CH_2_D_2_ values from cocultures are further away from isotopic equilibrium. From these experiments, we identify an energetic niche (ΔG_r_ ∼ −35 kJ mol^-1^ CH_4_), where *δ*^13^C, Δ^13^CH_3_D and Δ^12^CH_2_D_2_ values of hydrogenotrophic methanogenesis transiently exhibit more disequilibrium. This study presents differential responses of bulk carbon, hydrogen and clumped isotopic signatures to Δ G_r_, and highlights their applications in distinguishing methanogenesis under different energy regimes.

**ABSTRACT GRAPHICS:** 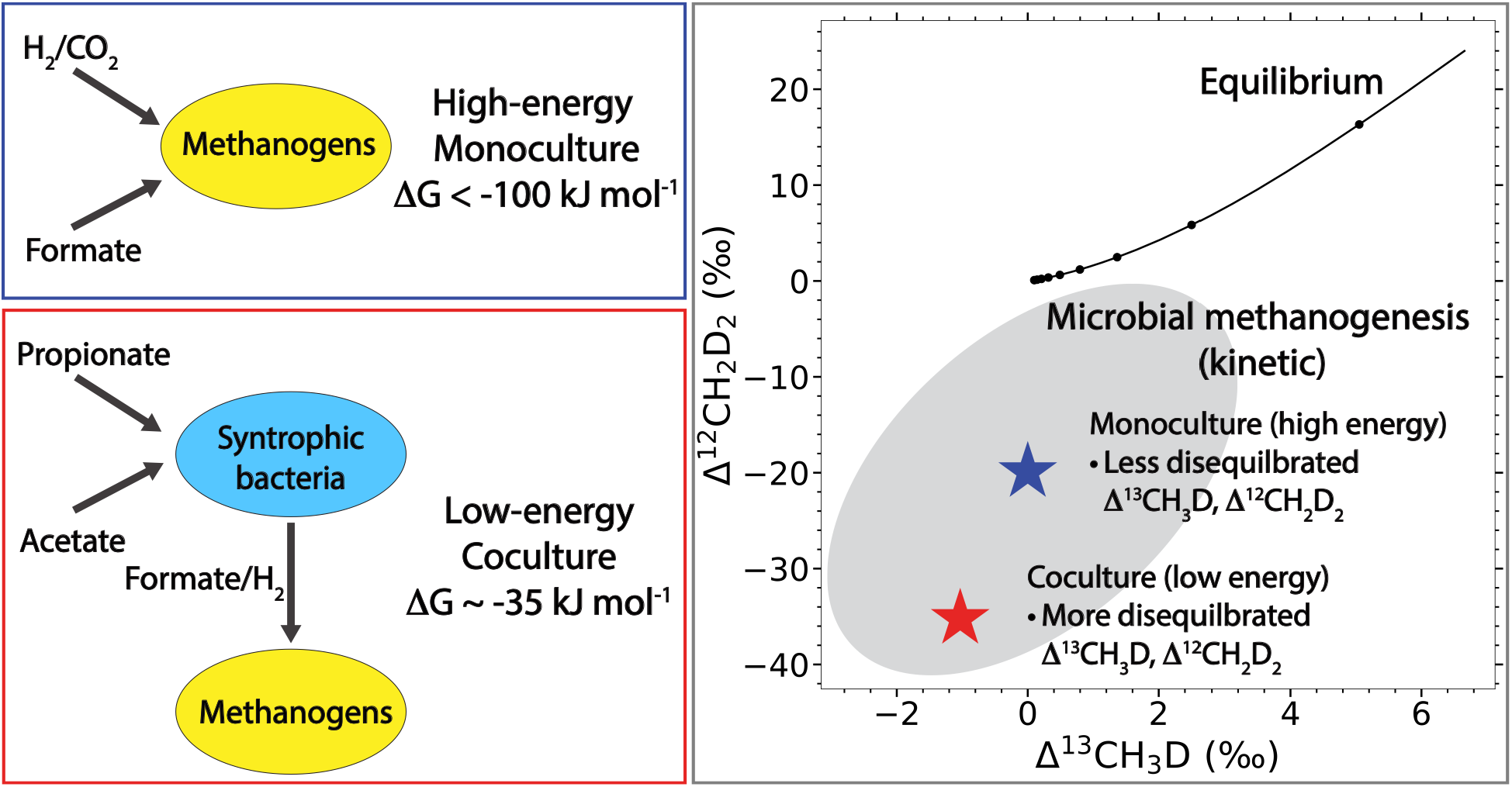

**SYNOPSIS:** This study demonstrates the compositions of doubly substituted isotopologues of methane as tracers for the thermodynamics of methanogenesis.

## Introduction

Methane is a potent greenhouse gas, contributing to ∼0.5 °C of warming between 2010 to 2019 relative to 1850 to 1900^1^, which is the most significant portion among all non-CO_2_ greenhouse gases. Due to the short residence time of methane, it is a good target for climate change mitigation^1,2^. Distinguishing and quantifying methane from different sources is therefore important for identifying major emission sources, improving estimates of the global methane budget, and evaluating targeted mitigation strategies. Reliable approaches for tracing methane sources could consequently contribute both to a better understanding of the global carbon cycle and to curbing the ongoing increase in atmospheric methane concentrations^2^.

Methane from microbial sources composes the majority of methane from wetlands, agriculture (e.g. livestock, rice paddies), landfills, and wastewater treatment systems, accounting for about two thirds of global emissions^3^. Methane is also the primary product of anaerobic digestion, a process widely used for biogas and biofertilizer production from organic waste streams^4^. Microbial methane is mostly produced via three biochemical pathways: hydrogenotrophic, methylotrophic and acetoclastic pathways^5,6^. Hydrogenotrophic methanogenesis uses H_2_ and CO_2_. In contrast, methylotrophic methanogenesis uses methylated organic compounds like methanol, methylamine and dimethyl sulfide, while acetoclastic methanogenesis uses acetate. In addition to directly using methylated organic compounds, methanogens can also pair with specific bacteria to perform syntrophic methanogenesis^7^. Syntrophic methanogenesis utilizes interspecies hydrogen or formate transport to couple the oxidation of organic molecules (e.g. acetate, propionate, butyrate) to methanogenesis^7,8^. During syntrophic methanogenesis, consumption of hydrogen by methanogenic archaea maintains low hydrogen partial pressure, thereby making the otherwise energetically unfavorable syntrophic oxidation reactions thermodynamically feasible^7,9^. This strategy sustains microbial activity close to the energy limit of life. Due to the low energy yield for the syntrophic microorganisms, this pathway is thermodynamically less competitive than acetoclastic methanogenesis under standard conditions. However, acetoclastic methanogens have lower resistance to stress occurring under thermophilic, acetate-limited or high-ammonia conditions, resulting in syntrophic methanogenesis becoming the dominant pathway^9–13^.

Identifying methanogenesis pathways in various environments is crucial to understanding the sources of methane emissions. Previous studies have attempted to use bulk hydrogen and carbon isotopes (*δ*^13^C and *δ*D) to trace microbial methanogenesis pathways in natural environments^14–16^. Yet, there are fundamental challenges where the isotopic signals of different sources and processes are overlapping in bulk isotope diagrams^16–18^. Advances in high-resolution mass spectrometry allow for the quantification of doubly substituted isotopologues of methane (Δ^13^CH_3_D and Δ^12^CH_2_D_2_, also called “clumped” isotopologues), which provides additional information on the formation processes of methane independently from the isotope values of source materials (e.g. water, CO_2_)^19–22^. Methane isotopic signatures can also vary with the energetics of methanogenesis due to reversibility of the methanogenic pathway catalyzed by methyl-coenzyme M reductase (Mcr)^23,24^. Methanogenesis proceeding with a high energy yield typically demonstrate non-equilibrium isotopic signatures as a result of low reaction reversibility^20,25^, while isotopic values move towards isotopic equilibrium as the available energy decreases and the reaction becomes more reversible^26,27^. These observations imply that isotopic signatures of methane are controlled by the combination of energetics and metabolic pathways under which methanogenesis occurs.

Despite previous studies examining the isotopic signatures of syntrophic methanogenesis under low energy yield^26,28^, several important knowledge gaps remain. Previous studies focused on the isotopic signatures of methanogenesis under high energy yield (ΔG < −90 kJ mol^-1^ CH_4_)^25,29^ and extremely low energy yield (ΔG > −30 kJ mol^-1^ CH_4_)^26,27^. How the clumped isotopic signatures of methane shift from high-energy to low-energy regimes are unclear. Second, the Δ^12^CH_2_D_2_ signatures of syntrophic methanogenesis are unknown, although it could potentially provide an important tracer for this process in natural and engineered environments. Third, the clumped isotopic signatures of formate-dependent methanogenesis remain unknown. However, formate can serve as an important interspecies electron carrier during the syntrophic degradation of organic acids, transferring reducing equivalents from syntrophic bacteria to methanogenic archaea. Characterizing the isotopic signatures of formate-dependent methanogenesis is therefore crucial to understanding the isotope signatures of syntrophic methanogenesis and completing the picture of biological source values.

To address these questions, we cultivated hydrogenotrophic methanogens in monoculture with abundant hydrogen or formate and compared their methane isotopic signatures with those produced under syntrophic conditions. For the latter, hydrogenotrophic methanogens were grown in coculture with hydrogen- and formate-producing syntrophic bacteria, where the energy availability is constrained for both microorganisms. We further investigated the effect of different substrates during the coculture cultivation, as the energy yield of hydrogen and formate production varies among different substrates (Table 1). We then mapped the variations in isotopic signatures of methane produced from different substrates over different energy regimes and compared these patterns with previous observations.

**Table 1.** Standard Gibbs energy changes of hydrogen-consuming methanogenesis and hydrogen-releasing reactions in monoculture and coculture experiments in this study.

| **Reaction** | **ΔG<sup>0'</sup> (kJ mol<sup>-1</sup>)<sup>†</sup>** |
|---|---|
| **Methanogenesis** |  |
| 4H<sub>2</sub> + CO<sub>2</sub> → CH<sub>4</sub> + 2H<sub>2</sub>O | –131.0 |
| 4HCOO<sup>-</sup> + H<sup>+</sup> + H<sub>2</sub>O → 3HCO<sub>3</sub><sup>-</sup> + CH<sub>4</sub> | –130.4 |
| **Hydrogen-releasing reaction** |  |
| CH<sub>3</sub>CH<sub>2</sub>COO<sup>-</sup> + 2H<sub>2</sub>O → CH<sub>3</sub>COO<sup>-</sup> + CO<sub>2</sub> + 3H<sub>2</sub> | 76.0 |
| CH<sub>3</sub>COO<sup>-</sup> + H<sup>+</sup> + 2H<sub>2</sub>O → 2CO<sub>2</sub> + 4H<sub>2</sub> | 94.9 |
| **Net reaction\*** |  |
| 4CH<sub>3</sub>CH<sub>2</sub>COO<sup>-</sup> + 2H<sub>2</sub>O → 3CH<sub>4</sub> + CO<sub>2</sub> + 4CH<sub>3</sub>COO<sup>-</sup> | –89.0 |
| CH<sub>3</sub>COO<sup>-</sup> + H<sup>+</sup> → CH<sub>4</sub> + CO<sub>2</sub> | –36.1 |

<sup>\*</sup> The hydrogen-releasing reactions of propionate and acetate are endergonic under standard states and thus require methanogenesis to drive them by lowering the hydrogen concentrations. After combining the methanogenesis and hydrogen-releasing half reactions, the net reactions are exergonic.
<sup>†</sup> The standard Gibbs energy data are from previous studies by Schink (1997) and Schauer et al. (1980)<sup>7,61</sup>.

## Materials and Methods

### Microbial cultivation

Cultivations of syntrophic cocultures on propionate were conducted at Dartmouth College with *Methanobacterium formicicum* (*M. formicicum*, DSM 1535, the methanogen) and *Syntrophobacter fumaroxidans* (*S. fumaroxidans*, DSM 10017, the syntrophic bacteria) obtained from DSMZ (Table 2). The C-P medium used in the incubation was adopted from previous studies^28,30^, with details and procedures in the Supplementary Materials (Text S1). Prior to the cultivation of cocultures, axenic pre-cultures of *M. formicicum* and *S. fumaroxidans* were grown in C-P medium separately until late exponential phase. *M. formicicum* was grown on 1.7 bar of 80%/20% (v/v) H_2_/CO_2_, while *S. fumaroxidans* was grown on 20 mM sodium fumarate (Text S1) with 1.7 bar of 80%/20% (v/v) N_2_/CO_2_ in the headspace. For the actual experiments, 47.5 mL of C-P medium were put into a 160-mL serum bottle and supplemented with 30 mM of sodium propionate. The medium was then inoculated with 1.5 mL of *M. formicicum* and 1.5 mL of *S. fumaroxidans* pre-cultures, respectively. The headspace was flushed with pure N_2_ and pressurized to 1.3 bar. The cocultures were then incubated statically at 37 and 30 °C. Two additional sets of experiment were done with the same amount of medium and culture. One set of experiments was conducted in 70-mL serum bottles under 37 °C, leaving a smaller headspace volume in each replicate. The other set, conducted at the University of Utah (UU), also used 160-mL serum bottles, but 80%/20% (v/v) Ar/CO_2_ instead of N_2_ was used as the headspace gas (Table 2). Each experiment was performed with five biological replicates. Cocultures were incubated for 52 to 85 days (Table S2). To compare between the coculture with monoculture, we also cultivated *M. formicicum* in monocultures in 160-mL serum bottles under the same conditions as the cocultures, except that only 2.5 mL of *M. formicicum* pre-cultures were used as inoculum, and the headspace was exchanged and pressurized with 1.7 bar of 80%/20% (v/v) H_2_/CO_2_ in substitution of propionate. The monoculture methanogen was incubated for 10 to 49 days (Table S2). The two microorganisms, *M. formicicum* and *S. fumaroxidans* are referred to as Mf and Sf, and the cocultures of them are referred to as Coc in the following text.

**Table 2.**
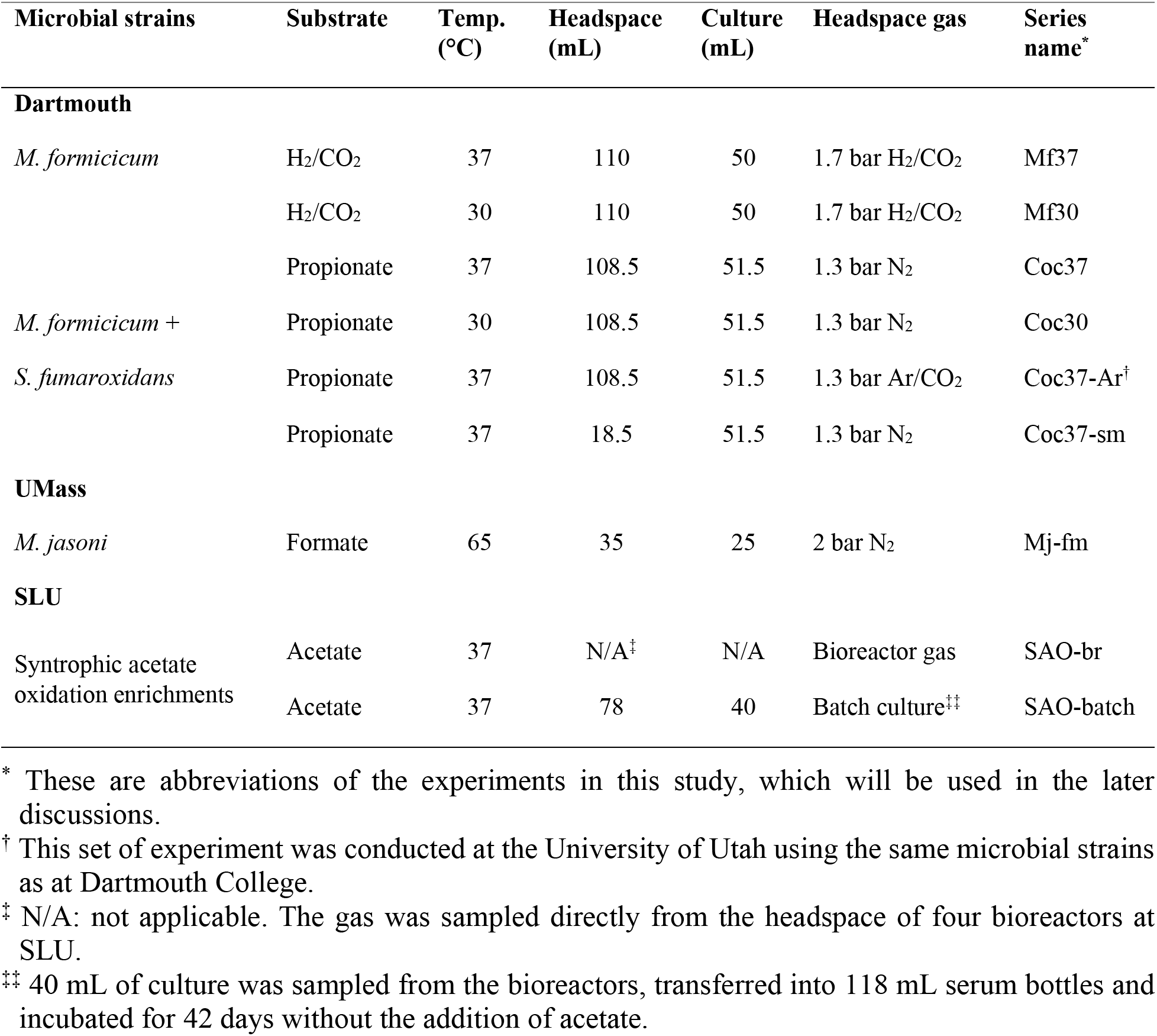
Experimental conditions for methanogenesis incubations.

| Microbial strains | Substrate | Temp. (°C) | Headspace (mL) | Culture (mL) | Headspace gas | Series name<sup>*</sup> |
|---|---|---|---|---|---|---|
| **Dartmouth** |  |  |  |  |  |  |
| *M. formicicum* | H<sub>2</sub>/CO<sub>2</sub> | 37 | 110 | 50 | 1.7 bar H<sub>2</sub>/CO<sub>2</sub> | Mf37 |
|  | H<sub>2</sub>/CO<sub>2</sub> | 30 | 110 | 50 | 1.7 bar H<sub>2</sub>/CO<sub>2</sub> | Mf30 |
| *M. formicicum* + *S. fumaroxidans* | Propionate | 37 | 108.5 | 51.5 | 1.3 bar N<sub>2</sub> | Coc37 |
|  | Propionate | 30 | 108.5 | 51.5 | 1.3 bar N<sub>2</sub> | Coc30 |
|  | Propionate | 37 | 108.5 | 51.5 | 1.3 bar Ar/CO<sub>2</sub> | Coc37-Ar<sup>†</sup> |
|  | Propionate | 37 | 18.5 | 51.5 | 1.3 bar N<sub>2</sub> | Coc37-sm |
| **UMass** |  |  |  |  |  |  |
| *M. jasoni* | Formate | 65 | 35 | 25 | 2 bar N<sub>2</sub> | Mj-fm |
| **SLU** |  |  |  |  |  |  |
| Syntrophic acetate oxidation enrichments | Acetate | 37 | N/A<sup>‡</sup> | N/A | Bioreactor gas | SAO-br |
|  | Acetate | 37 | 78 | 40 | Batch culture<sup>‡‡</sup> | SAO-batch |

<sup>\*</sup> These are abbreviations of the experiments in this study, which will be used in the later discussions.
<sup>†</sup> This set of experiment was conducted at the University of Utah using the same microbial strains as at Dartmouth College.
<sup>‡</sup> N/A: not applicable. The gas was sampled directly from the headspace of four bioreactors at SLU.
<sup>‡‡</sup> 40 mL of culture was sampled from the bioreactors, transferred into 118 mL serum bottles and incubated for 42 days without the addition of acetate.

*Methanothermococcus jasoni* (*M. jasoni*, referred to as Mj) was cultivated at the University of Massachusetts, Amherst (UMass) as previously described^31^. Briefly, 25 mL of DSM 282 medium (Text S2) with 0.1% (w/v) sodium formate was transferred into butyl rubber stopper sealed 60-mL serum bottles. The medium was degassed and flushed with 2 bar (70 mL) of N_2_ gas and reduced with 0.025% (w/v) each of cysteine-HCl and Na_2_S·9H_2_O. Three biological replicates were inoculated with 0.1% (v/v) of Mj at exponential growth phase, then incubated at 65 °C for 9 to 12 hours (Table S3).

The growth conditions of the four bioreactors at the Swedish University of Agricultural Sciences (SLU) were described in a previous study^32^, with the modification that all of the bioreactors were fed with anaerobic medium containing 100 mM sodium acetate as the sole substrate at the time of sampling. The bioreactors were operated at 37 °C with the same inflow rate of growth medium, giving a hydraulic retention time (HRT) of 28 days. Total ammoniacal nitrogen (TAN) concentrations and pH were different among bioreactors, leading to different free ammonia nitrogen (FAN), as shown in Table S1. The acetate supplied to the bioreactors was predominantly converted to methane via syntrophic acetate oxidation (SAO), as supported by 16S rRNA gene sequencing of the enrichment cultures, showing low (< 0.5% in relative abundance of the microbial community) or no presence of acetoclastic methanogens (Figure S1). The 16S rRNA sequencing and data analysis follow the method described in previous studies^32,33^. Headspace gas of each bioreactor was sampled by connecting an evacuated 118-mL serum bottles sealed with black butyl septa and equilibrated for 10 minutes. Additionally, the enrichment cultures of the bioreactors were sampled and cultivated in batch. Four anoxic bottles (118 mL) were prepared by flushing with pure N_2_ and exchanging the gas three times with N_2_/CO_2_ and were autoclave-sterilized. Then 40 mL of enrichment culture were directly transferred from the bioreactors to the anaerobic bottles using a syringe, followed by static incubations in dark at 37 °C for 42 days without adding extra acetate.

### Headspace gas concentration measurement

Gas concentrations in the headspace were measured by a dual-channel gas chromatography (GC, SRI 8610C) with a flame ionization detector (FID) and a thermal conductivity detector (TCD). The GC is equipped with a 6’ x 1/8” column filled with HayeSep D polymer. Helium (UHP, AirGas) was used as the carrier gas during the measurements of methane, carbon dioxide, and normal amounts of hydrogen (for incubations with H_2_/CO_2_ as the headspace gas). For the incubation of cocultures, trace amounts of H_2_ were below the detection limit on TCD with helium as the carrier gas. Therefore, argon (UHP, Airgas) was used as carrier gas for the quantification of trace hydrogen. Peak areas of the gas on the GC were integrated with the PeakSimple software. A calibration with 1% (v/v) methane/helium standard (AirGas) was done to ensure the analytical linearity of the instrument prior to each daily measurement of methane. For normal amounts of hydrogen and carbon dioxide, 80%/20% (v/v) H_2_/CO_2_ gas (Airgas) was used as the standard. For trace amount of hydrogen gas, the standard samples were made by diluting 95%/5% (v/v) N_2_/H_2_ gas (Airgas) in air. Four mixtures were made with different H_2_ concentrations at 50, 100, 199 and 331 ppm, respectively.

Depending on headspace gas concentrations, 5 to 1000 *μ*L of gas was sampled from each bottle using calibrated gas-tight syringes with valves (Hamilton Company). The uncertainties of measurements on methane, carbon dioxide and normal hydrogen gas concentrations were reported as the 1 standard deviation among biological replicates for the experiments at Dartmouth. For the experiments at SLU, the 1 standard deviation of 3 repeated measurements on the same sample was reported as the uncertainty (Table S2), since the samples came from the headspace of individual bioreactors. The uncertainties of the measurements on trace amount of hydrogen gas in coculture cultivation experiments at Dartmouth, UU and SLU were propagated from the uncertainties of the slope and intercept of the calibration curves.

### Stable Isotope notations

The total ^13^C/^12^C and D/H ratios (bulk isotope ratios) are expressed by the *δ* notations in per mil (‰) relative to international standard materials (VPDB and VSMOW):

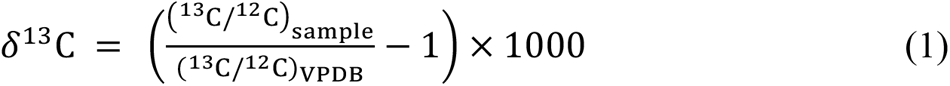

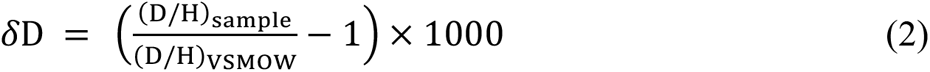

The clumped isotopologues compositions are expressed in Δ notation relative to the stochastic state, where carbon and hydrogen isotopes are randomly distributed among methane isotopologues^34^:

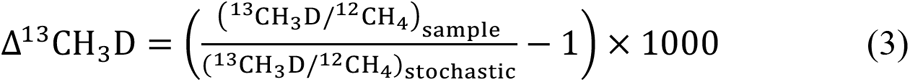

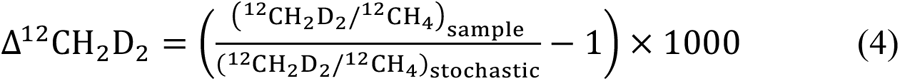

The temperature dependence of Δ^13^CH_3_D and Δ^12^CH_2_D_2_ at intramolecular thermodynamic equilibrium is adopted from the *ab initio* calculations^21^.

Isotopic fractionations between methane and substrate (water or carbon dioxide) are expressed by fractionation factors *α* and ε. The fractionation factor *α* denotes the relative isotopic ratios in two reservoirs:

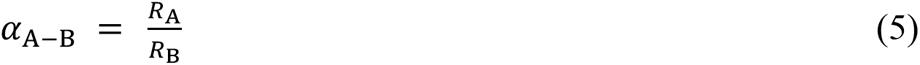

Where *R*_A_ and *R*_B_ are the isotopic ratios (^13^C/^12^C or D/H) in reservoir A and B, respectively. To compare with the data in previous studies, *α*_A-B_ is converted into *ε*_A-B_ in per mil (‰):

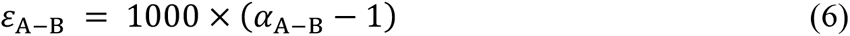

### Stable Isotope measurements

The isotopic signatures of methane were measured on the Nu instrument gas-source high-resolution isotope ratio mass spectrometer (Panorama) at UCLA^35^. The purification and analytical procedures of methane followed the protocols described in the previous studies^25,36,37^. Due to the large amounts of N_2_ relative to CH_4_ in the headspace gas of coculture experiments, the purification procedures were conducted twice to ensure a complete separation of CH_4_ from N_2_.

For the analysis of a gas sample, *δ*^13^C and *δ*^13^CH_3_D were measured by 10 to 20 blocks of 20 pairs of sample-reference gas measurement cycles, while *δ*D and *δ*^12^CH_2_D_2_ were measured with 28 to 40 blocks of sample-reference measurement cycles. Smaller samples were measured by fewer gas measurement cycles and typically had larger uncertainties (as shown in Table S3). The external precision of the isotopic measurements was evaluated by repeated measurements of a Utica gas sample (a thermogenic methane gas standard at UCLA). The average and standard deviations (2*σ*) of 26 measurements from 2016 to 2026 were −25.656 ± 0.323 ‰, −153.989 ± 0.679 ‰, 3.27 ± 1.23 ‰ and 8.94 ± 1.98 ‰ for *δ*^13^C, *δ*D, Δ^13^CH_3_D and Δ^12^CH_2_D_2_, respectively. The hydrogen isotope (*δ*D) of water used in the cultivation at Dartmouth was measured at Dartmouth Stable Isotope Laboratory (Table S4) via the method described in a previous study^38^. The results were reported as *δ*D relative to VSMOW on the VSMOW-SLAP scale with 1*σ* uncertainties smaller than 0.5 ‰. The hydrogen isotope of water from the cultivation at Utah, UMass and SLU was measured at Stable Isotope Ratio Facility for Environmental Research (SIRFER) at the University of Utah (Table S4), following the method described elsewhere^39^. The uncertainties of the measurements were taken as the 1 standard deviation of 5 repeated measurements on the reference material used at SIRFER.

The carbon isotope (*δ*^13^C) of carbon dioxide was measured at the Stable Isotope Facility (SIF) at UC Davis, using a ThermoScientific GasBench system interfaced to a ThermoScientific Delta V Plus isotope ratio mass spectrometer (ThermoScientific, Bremen, Germany). CO_2_ was first sampled by a six-port rotary valve (Valco, Houston TX) in the helium carrier gas and then separated from other residual gases by a 25 m × 0.32 mm (inner diameter) Poraplot Q GC column at 45 °C and with 2.5 mL/min of helium flow. A pure reference gas (CO_2_) was used to calculate provisional delta values of the sample peak. The final *δ*^13^C values were obtained after adjusting the provisional values for changes in linearity and instrumental drift. At least two laboratory reference materials (calibrated directly against NIST 8545) were analyzed with every 10 samples. The final *δ*^13^C values were expressed relative to VPDB.

### Energy yield calculations

To investigate the variation of methane isotope signatures in response to Gibbs energy availability, we estimated the ΔG_r_ values of our experiments using the following equation:

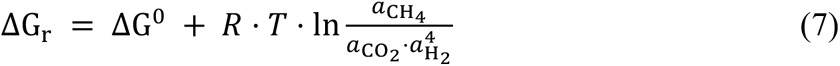

Where ΔG^O^ is the standard molar Gibbs energy at the given temperature, calculated from the Van’t Hoff equation^40^, and *a*_CH4_, *a*_CO2_ and *a*_H2_ are the activities of dissolved gas molecules, which were calculated from the partial pressure of the gas and the corrected solubility (in mol L^-1^ bar^-1^). The corrected solubility of methane and carbon dioxide was determined by the Setschenow equation, using the Setschenow constant (*K*_salt_), the molar concentration of electrolyte (*C*_salt_), and the standard solubility (*S*_O_) in pure water at 1 bar^41^:

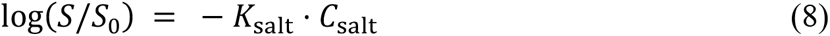

The concentration of electrolyte in the medium was estimated to be 0.1 mol L^-1^. The Setschenow constants of the gases^42,43^ and standard solubility^44,45^ from the previous studies were used to calculate the corrected solubility for methane and carbon dioxide. The solubility of hydrogen gas shows little variation in the range of medium salinity in this study^46^ and thus the standard solubility was used.

### Isotope modeling

The variations of isotopic signatures with ΔG_r_ of hydrogenotrophic methanogenesis were calculated from the isotopic model developed in the previous study^24^. Briefly, the reversibility of hydrogenotrophic methanogenesis was constrained by the concentrations of dissolved hydrogen, inorganic carbon, and methane. The net bulk and clumped isotope effects were calculated by the reversibility and kinetic isotope effect (KIE) of each step^24^. Due to the uncertainties in KIE at each step of hydrogenotrophic methanogenesis, 200 sets of KIE values were drawn randomly from the distribution of each KIE in the model^24^, with a combination of 14 KIEs for hydrogen and 8 KIEs for carbon in each set. For consistency with our experiments, we set the hydrogen and carbon isotope values of water and carbon dioxide to −53 and −7.5 ‰, respectively, and temperature was set to 35 °C. These values are close to the incubation temperature and the measured isotope values of water and carbon dioxide used by the cultivation of Mf+Sf coculture (Table S4). The median, 68 % and 95 % of the resulting isotopic signatures and fractionations were calculated and presented as the result.

## Results

Methane production rate is generally higher for monoculture methanogens grown on abundant H_2_/CO_2_ or formate compared with that for coculture grown on propionate. The change of headspace gas compositions in monoculture (Mf) and coculture (Mf+Sf) cultivations is plotted on Figure S2. At the optimal growth temperature (37 °C), methane production was 54 times faster in the Mf monoculture supplied with H_2_/CO_2_ than in the syntrophic Mf+Sf coculture grown on propionate, with methane production rates of 0.867 ± 0.040 *μ*mol mL^-1^ day^-1^ and 0.016 ± 0.001 *μ*mol mL^-1^ day^-1^, respectively (Table S5). Lowering temperature to 30 °C decreased methane production rate of the Mf monoculture to 0.339 ± 0.015 *μ*mol mL^-1^ day^-1^. In contrast, the Mf+Sf cocultures at 30 °C produced methane at 0.016 ± 0.003 *μ*mol mL^-1^ day^-1^, which is similar to the rate for coculture at 37 °C. Compared to Mf grown on H_2_/CO_2_, methane production rate of Mj grown on formate was an order of magnitude higher (Table S5). The Mf monoculture consumed 85 to 89 % of initial H_2_ at the end of the incubation (Figure S2, Text S3). On the other hand, due to the slow methane production by the syntrophic Mf+Sf coculture, it is estimated that only 3 to 10 % of the initial propionate was consumed at the end of the cultivation (Text S3).

The Gibbs energy, ΔG_r_, of methanogenesis by the Mf monocultures ranged from −127.9 ± 0.8 to −101.7 ± 0.8 kJ mol^-1^ CH_4_ at 37 °C (Table S2). Monoculture methanogenesis at 30 °C showed a similar ΔG_r_ as the last time point of measurements at 37 °C, with ΔG_r_ = −108.3 ± 1.3 kJ mol^-1^ CH_4_ (Table S2). In monoculture methanogenesis experiments, ΔG_r_ became less negative as the experiments proceed due to the consumption of substrates (H_2_ and CO_2_) and production of methane. In contrast, substantially less energy was available from methanogenesis under syntrophic conditions. The Gibbs energy yield ranged from −41.0 ± 1.4 to −36.5 ± 1.5 kJ mol^-1^ CH_4_ for the Mf+Sf cocultures and from −39.6 ± 0.6 to −25.6 ± 2.3 kJ mol^-1^ CH_4_ for the bioreactors with SAO enrichments (Table S2).

The results demonstrated that methane produced from mono-versus cocultures could be differentiated by unique isotopic compositions (Figure 1). In general, *δ*^13^C values were more negative for methane produced by cocultures, ranging from −94.332 ± 0.014 to −85.528 ± 0.005 (Coc and SAO series on Figure 1A, Table S3), whereas hydrogenotrophic methanogenesis by monoculture Mf exhibited less negative *δ*^13^C values from −58.661 ± 0.040 to −56.395 ± 0.004 ‰ (Mf series on Figure 1A, Table S3). Monoculture methanogen grown on formate yielded methane *δ*^13^C values between monoculture and coculture hydrogenotrophic methanogenesis (Mj series on Figure 1A, Table S3). Methane *δ*D values, on the other hand, did not show distinct patterns among different methane production pathways (Figure 1), with a range from −405.168 ± 0.041 to −371.024 ± 0.023 ‰ (Table S3). In terms of clumped isotopologues compositions of methane, monoculture methanogenesis on formate and H_2_/CO_2_ (Mf and Mj series on Figure 1B) generally produced less negative Δ^13^CH_3_D and Δ^12^CH_2_D_2_ values than coculture methanogenesis on acetate and propionate (Coc and SAO series on Figure 1B). The former possesses Δ^13^CH_3_D values from −1.14 ± 0.14 to 2.61 ± 0.24 ‰ and Δ^12^CH_2_D_2_ values from −27.04 ± 0.80 to −20.64 ± 0.90 ‰, while the latter yielded Δ^13^CH_3_D values from −2.32 ± 0.37 to 0.44 ± 0.19 ‰ and Δ^12^CH_2_D_2_ values from −38.46 ± 0.89 to −30.56 ± 1.42 ‰, respectively (Table S3).

**Figure 1.**
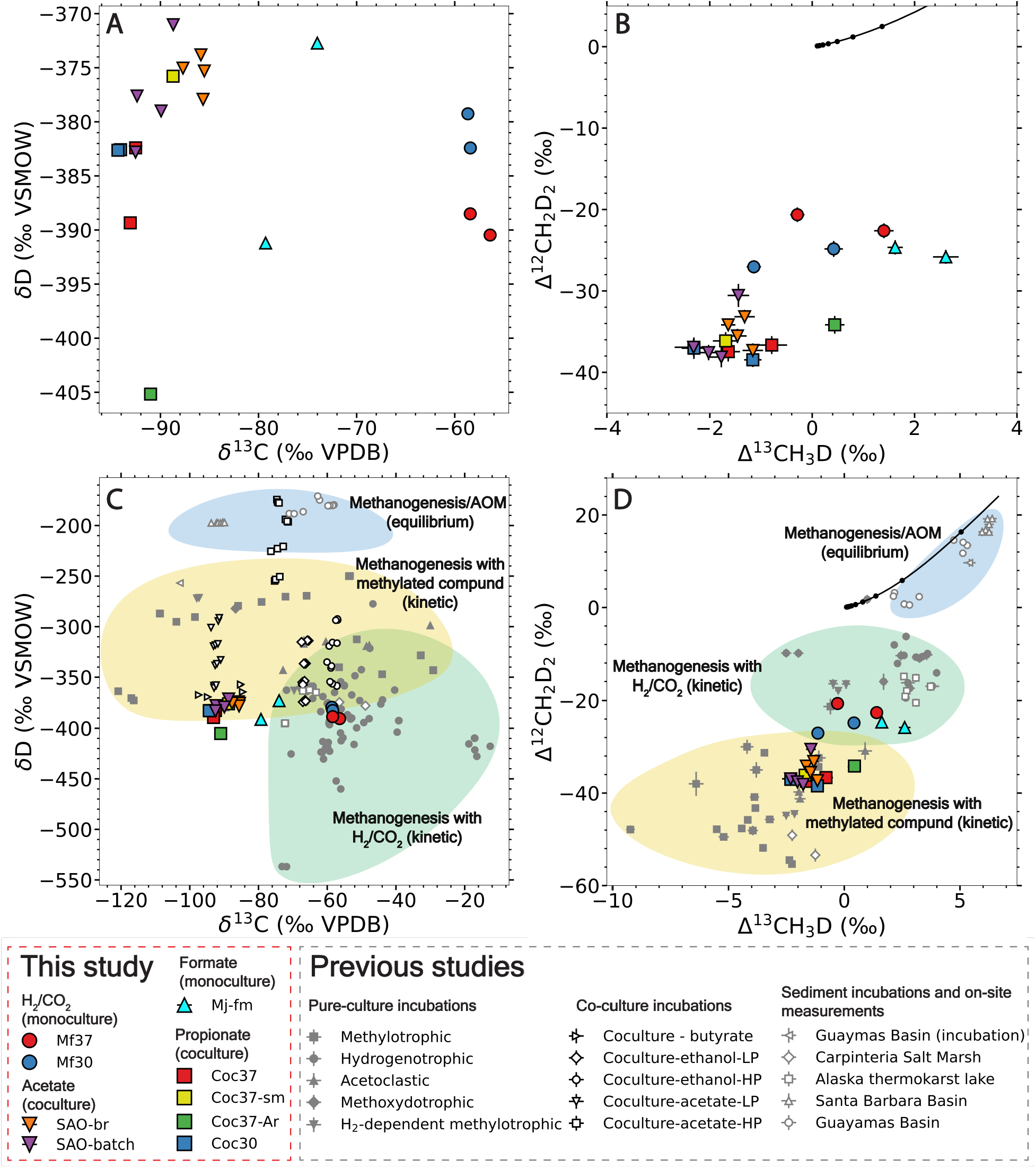
Isotopic signatures of methane in monoculture and coculture cultivations in (A) bulk isotope (*δ*D – *δ*^13^C) and (B) clumped isotope (Δ^12^CH_2_D_2_ – Δ^13^CH_3_D) spaces, and comparisons of the isotopic signatures with previous studies in (C) bulk and (D) clumped isotope spaces. The shortened names of each experiment in the legend can be found in Table 2. Data from previous studies are shown as points in gray or white on panels C and D, and encompass pure-culture methanogenesis incubations via different pathways^21,26,29,47,59,65,73,74^, coculture methanogenesis incubations on different substrates^26,65^, and sediment incubations and on-site measurements^27^. The “HP” and “LP” in the legend denotes incubations under high hydrostatic and atmospheric pressure^26^. The yellow, green and blue-shaded areas are regimes of kinetic methanogensis with methylated compounds, with H_2_/CO_2_ and equilibrium-driven methanogenesis and AOM that are collectively defined by these studies.

Isotopic fractionations of hydrogen and carbon between methane, water and carbon dioxide (^D^*ε*_CH4-H2O_ and ^13C^*ε*_CH4-CO2_) tracked the pattern of bulk *δ*^13^C and *δ*D of methane, as shown in Figure 2 and Table S4. Hydrogen isotope fractionation factor between methane and water (^D^ε_CH4-H2O_) ranged from −356.45 ± 0.11 to −325.67 ± 0.15 ‰. Carbon isotope fractionation (^13*C*^ε_CH4-CO2_) was the largest in the coculture methanogenesis (−87.44 ± 0.07 to −78.60 ± 0.07 ‰), followed by monoculture methanogenesis with formate (−66.78 ± 0.07 and −72.09 ± 0.07 ‰). The smallest carbon isotope fractionation was observed in monoculture methanogenesis grown on H_2_/CO_2_ (−59.13 ± 0.07 to −40.86 ± 0.07 ‰). Results from the isotopic model adopted from the previous study by Gropp et al.^24^ are shown in Figure 3.

**Figure 2.**
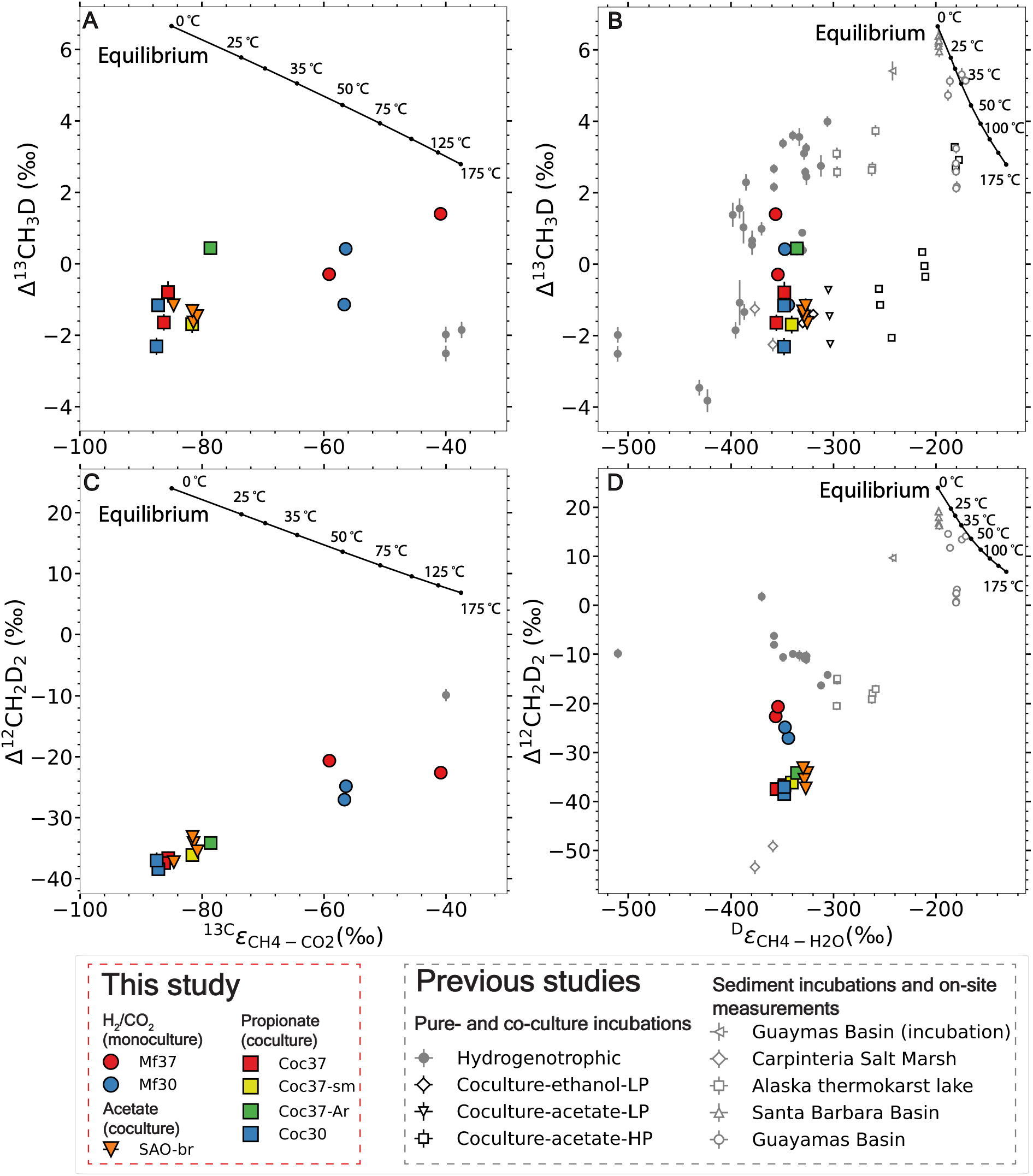
Variations in isotopic fractionation with clumped isotopic signatures in (A) Δ^13^CH_3_D − ^13C^*ε*_CH4-CO2_; (B) Δ^13^CH_3_D − ^D^*ε*_CH4-H2O_; (C) Δ^12^CH_2_D_2_ − ^13C^*ε*_CH4-CO2_ and (D) Δ^12^CH_2_D_2_−^D^*ε*_CH4-H2O_ spaces. Previous data from microbial incubations and on-site measurements are included for comparison^25–27,59^. The “HP” and “LP” in the legend denote incubations under high hydrostatic and atmospheric pressure, respectively^26^. The line in black shows the isotopic equilibrium of bulk carbon isotope between CH_4_ and CO_2_^75^ and bulk hydrogen isotope between CH_4_ and H_2_O^76^, as well as intramolecular isotopic equilibrium values of two clumped isotopologues of methane^21^ from 0 to 175 °C. The values of hydrogen and carbon isotope fractionation factors are listed in Table S6.

**Figure 3.**
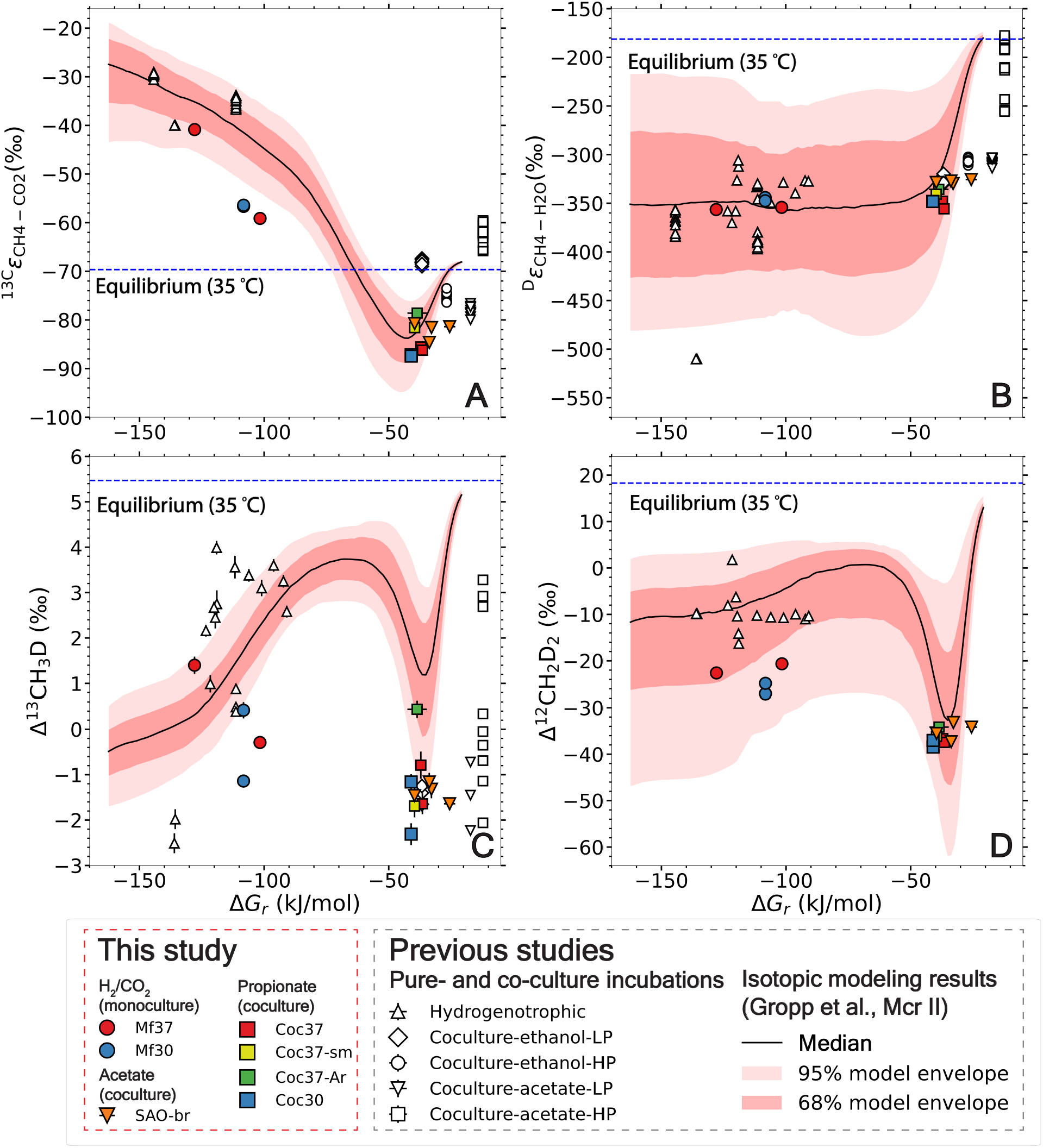
Variations of isotope fractionation and isotopic signatures with the ΔG_r_ of hydrogenotrophic methanogenesis: (A) ^13C^*ε*_CH4-CO2_; (B) ^D^*ε*_CH4-H2O_; (C) Δ^13^CH_3_D and (D) Δ^12^CH_2_D_2_. For comparison, data from previous monoculture and coculture incubations^25,26^ are presented as points in white. The median from 200 simulations by previous isotopic model with the key enzyme Mcr II^24^ are shown as black lines, and the areas in red and light red represent the envelopes encompassing 68 % and 95 % of the results. The dashed line in blue denotes the equilibrium values at 35 °C^21,75,76^.

## Discussion

### Isotopic signatures of methane from monocultures using H_2_/CO_2_ and formate

Methane isotopic signatures of monocultures grown on H_2_/CO_2_ largely overlap with the kinetic hydrogenotrophic methanogenesis space defined by previous studies (Figure 1). Notably, the Δ^12^CH_2_D_2_ values of methane produced by monoculture *M. formicicum* are more negative than previous observations of hydrogenotrophic methanogenesis with strains from genus *Methanosarcina*, *Methanocaldococcus*, and *Methanococcus*, which typically range between −9 to −20 ‰^21,25,47^. The ΔG_r_ of hydrogenotrophic methanogenesis in the previous study is estimated to range from −140 to −100 kJ mol^-1^ CH_4_^25^, similar to this study. Therefore, the Gibbs energy of methanogenesis is unlikely to be the reason for different Δ^12^CH_2_D_2_ values. Clumped isotope signatures of methane produced by methanogens from *Methanosarcina* and *Methanococcus*/*Methanocaldococcus* have been reported^25^, while this is the first report of methane clumped isotopic signatures produced by the genus *Methanobacterium*. It has been shown that during methanogenesis, the combination of hydrogen from sources with different hydrogen isotope values can generate various Δ^12^CH_2_D_2_ values of methane^25,36,48^. This “combinatorial effect” causes more negative Δ^12^CH_2_D_2_ values when the intermediates of methanogenesis and intracellular hydrogen carriers have more different hydrogen isotope values^49^. It is plausible that *Methanobacterium formicicum* has genus-specific D/H fractionations that is different from other methanogens, which yields larger differences between the δD values of intermediates and hydrogen carriers and thus more negative Δ^12^CH_2_D_2_ values.

During microbial methanogenesis with formate, the substrate is first oxidized to CO_2_ by formate dehydrogenase (Fdh)^50^ and coenzyme F_420_ is reduced to F_420_H_2_, which is further used for the reduction of CO_2_ to methane^51^. Although trace H_2_ is produced during the decomposition of formate, it has been proposed to not directly contribute to methanogenesis^51^. Therefore, methanogenesis with formate in principle follows the same pathway as hydrogenotrophic methanogenesis but utilizing the reducing power from formate rather than H_2_. Methanogenesis from formate in the present study indeed exhibited similar isotopic values to monoculture methanogenesis from H_2_/CO_2_ in *δ*D, Δ^13^CH_3_D and Δ^12^CH_2_D_2_. However, the *δ*^13^C values are more negative than the typical hydrogenotrophic methanogenesis under high Gibbs energy yield (Figure 1A and 1C). Meanwhile, the Δ^12^CH_2_D_2_ values are also more negative compared with hydrogenotrophic methanogenesis by the genus *Methanococcus* and *Methanocaldococcus*^25^ (Figure 1B and 1D). Overall, methanogenesis from formate by Mj demonstrates a shift in *δ*^13^C and Δ^12^CH_2_D_2_ values towards those of syntrophic methanogenesis from propionate and acetate (Figure 1). This suggests that the formate-consuming methanogens in this study perform hydrogenotrophic methanogenesis at a lower Gibbs energy yield, possibly utilizing the trace hydrogen derived from formate.

### Isotopic values of hydrogenotrophic methanogenesis reflect a thermodynamic niche

The isotope signatures of methane produced from syntrophic propionate and acetate oxidation generally demonstrate more negative *δ*^13^C, Δ^13^CH_3_D and Δ^12^CH_2_D_2_ values compared with monoculture methanogenesis from H_2_/CO_2_ and formate (Figure 1), reflecting variations in the isotope signatures of hydrogenotrophic methanogenesis with the energy yield. The Gibbs energy yield (ΔG_r_) per methane molecule produced by methanogens in syntrophic cocultures is less than the monoculture under H_2_-rich conditions (Table S2). This results from the very low H_2_ concentrations required to make syntrophic oxidation of organic acids thermodynamically feasible, which simultaneously decreases the thermodynamic drive for hydrogenotrophic methanogenesis^7,8^. Less negative ΔG_r_ leads to more reversibility of the same enzymatic reaction^52^, and correspondingly the net isotope effect is expected to be closer to the equilibrium isotope effect (EIE). However, this seems to contradict with the experimental observations of methane isotopic signatures, where both Δ^13^CH_3_D and Δ^12^CH_2_D_2_ are further away from isotopic equilibrium (Figure 2 and 3). Previous studies also report a deviation from isotopic equilibrium of Δ^13^CH_3_D during methanogenesis under energy-limited conditions^26,53^. This study provides evidence that Δ^12^CH_2_D_2_ values have a finite deviation from isotopic equilibrium under energy-limited conditions as well, which is predicted by some earlier models^24,53^. Specifically, the experimental results in this study fit the trend of the isotopic model developed by Gropp et al.^24^ (Figure 3, Table S5). This deviation of clumped isotopic signatures when ΔG_r_ is around −35 kJ mol^-1^ is attributed to a differential stepwise response of the reaction reversibility to the change of ΔG_r_ (Figure 4). The Mcr- and Mtr-catalyzed steps for methane assembly stay largely irreversible when ΔG_r_ is more negative than −35 kJ mol^-1^, while the reversibility of the other steps of methanogenesis gradually decreases with more negative ΔG_r_ in this range^24^. The consistent expression of the KIE of Mcr- and Mtr-catalyzed steps due to the low reversibility results in the disequilibrium isotope signals in both mono- and cocultures, and the difference between these two originates from a shift from EIE towards KIE at the upstream steps of methanogenesis with more negative ΔG_r_^24^ (Figure 4).

**Figure 4.**
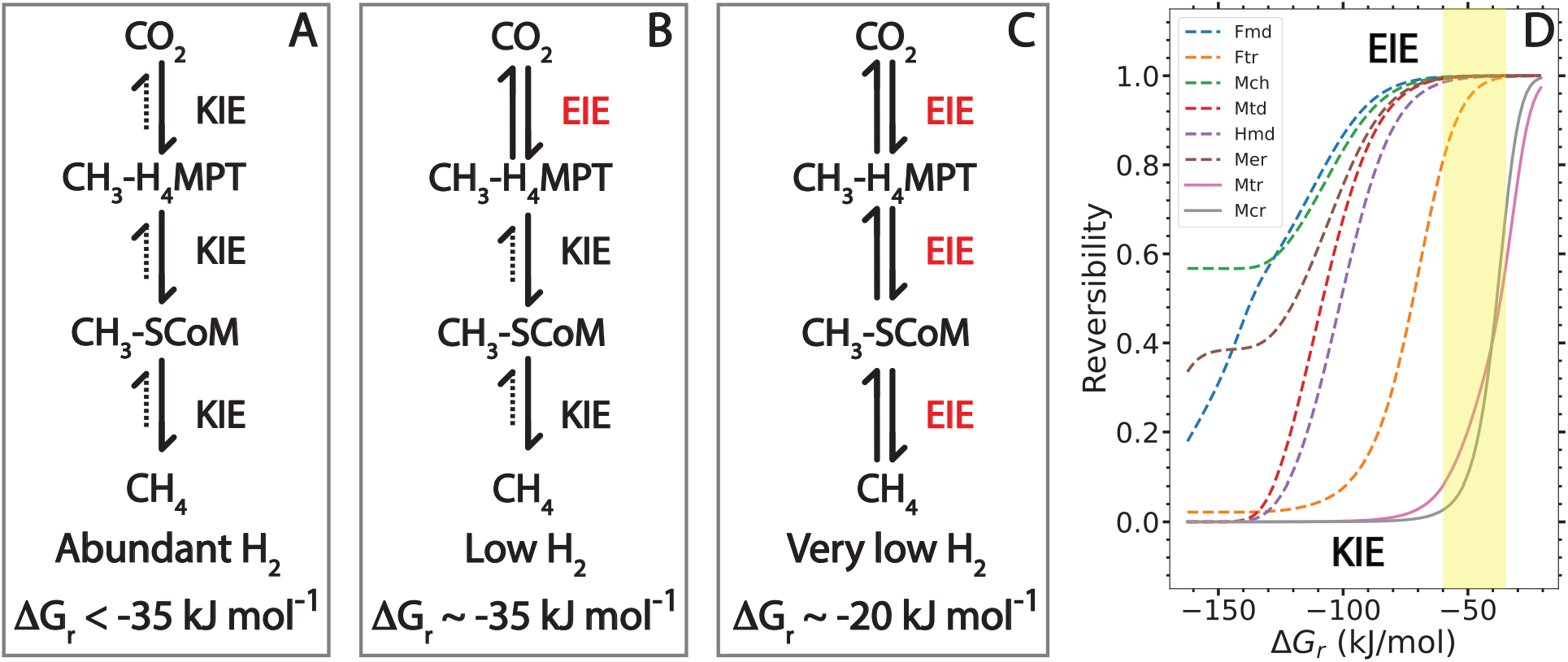
An illustration of the mechanism for the variation in methane isotopic signatures with (A) abundant H_2_ and high Gibbs energy yield (< −35 kJ mol^-1^); (B) low H_2_ and a Gibbs energy yield around the ‘energy niche’ at −35 kJ mol^-1^; and (C) very low H_2_ and a Gibbs energy yield close to the energy limit of microbial metabolism (∼ −20 kJ mol^-1^). Panel D shows the change in reversibility of each enzymatic reactions with the ΔG_r_ of methanogenesis in the model^24^. The intermediates of methanogenesis are noted as CH_3_-H_4_MPT (methyl-tetrahydromethanopterin) and CH_3_-SCoM (methyl-coenzyme M). The reactions between CO_2_ and CH_3_-H_4_MPT are catalyzed by a series of enzymes including Fmd (formylmethanofuran dehydrogenase), Ftr (formylmethanofurantetrahydromethanopterin formyl-transferase), Mch (methenyltetrahydromethanopterin cyclohydrolase), Mtd (methylenetetrahydromethanopterin reductase), Hmd (H_2_-forming methylenetetrahydromethanopterin dehydrogenase) and Mer (5,10-methylenetetrahydromethanopterin reductase)^24^. The last two steps of methanogenesis from CH_3_-H_4_MPT to CH_4_ are catalyzed by Mtr (tetrahydromethanopterin S-methyl-transferase) and Mcr (methyl-coenzyme M reductase)^77^. The dashed arrows on panels A-C represent smaller fluxes in the backward direction relative to the forward direction. This results in a lower reaction reversibility (closer to 0) and an isotope effect that is closer to the kinetic isotope effect (KIE). Conversely, solid arrows in both directions represent a higher reaction reversibility (closer to 1) and an isotope effect that is closer to the equilibrium isotope effect (EIE). The yellow area on panel D marks the range of ΔG_r_ between −60 to −35 kJ mol^-1^, where Δ^13^CH_3_D and Δ^12^CH_2_D_2_ values get further away from equilibrium (Figure 3C and 3D). This corresponds to high reversibility of the upstream steps and relatively lower reversibility of the last two steps.

The low reversibility of the Mcr-catalyzed steps across high- and low-energy methanogenesis in this study likely reflects an important regulation on the expression of the two Mcr isoenzymes - Mcr I and II - by the methanogens^54^. When the methanogens have replete H_2_, the low-affinity Mcr II is synthesized, while the high-affinity Mcr I is synthesized when H_2_ is limited^54–56^. This regulation ensures low reversibility of the Mcr-catalyzed step and net methane production even at low ambient H_2_ concentrations. However, the isotopic model using Mcr I points to a more negative ΔG_r_ value (∼−45 kJ mol^-1^) at the ‘energy niche’ with the largest degree of disequilibrium, showing a discrepancy from the experimental observations (Figure S3). A plausible explanation is that Gibbs energy yield of the experiments in this study may be underestimated due to inter-species electron transfer, which increases the effective hydrogen availability for the methanogen than that calculated from the partial pressure of hydrogen^8^. Especially when cells form aggregates, the inter-species hydrogen transfer is greatly enhanced^57^. Therefore, the actual ΔG_r_ of the methanogenesis reactions in this study may be more negative than the estimation based on measured H_2_, CO_2_ and CH_4_ concentrations. Despite the uncertainties in the exact values of ΔG_r_, methanogenesis by syntrophic cocultures generally yields much less energy than by monocultures.

Although the clumped isotope signatures of methane produced by cocultures are similar to those reported for acetoclastic and methylotrophic methanogenesis (Figure 1D)^25,58,59^, hydrogenotrophic methanogenesis is likely to be the dominant methanogenesis pathway in the coculture cultivations for several reasons. First, the methanogen used in the cultivation of propionate-fed coculture (Mf) can only use H_2_/CO_2_ or formate for methanogenesis^60,61^. In terms of the acetate-fed enrichments (SAO enrichments) in this study, they contain high levels of syntrophic acetate oxidizing bacteria and hydrogenotrophic methanogens, but no acetoclastic methanogens, except in reactor SL1 with lower TAN and FAN (Figure S1), where acetoclastic methanogens are present at very low levels (< 0.5%). The very negative *δ*^13^C values of methane are also distinct from the typical range of acetoclastic methanogenesis (−70 to −45 ‰)^16,25,58,59^. Collectively, these suggest hydrogenotrophic methanogenesis as the dominant pathway in the experiments of this study.

Formate can also serve as an inter-species electron carrier between syntrophs^8,9^. Due to the lack of constraints, however, the proportion of formate-relative to hydrogen-based electron transfer cannot be determined in this study. The similarity of isotopic signatures during methanogenesis with H_2_/CO_2_ and formate indicates that it is unlikely the main driver of the isotopic difference between monoculture and coculture methanogenesis (Figure 1). As such, the energetics of methanogenesis remains the most plausible driver for the observed trend in the isotopic signatures in this study.

The experimental and modeling results in this study reveal differential responses of carbon, hydrogen, and clumped isotopologue fractionations to Gibbs energy yield (Figure 3). When ΔG_r_ becomes less negative from −150 to −35 kJ mol^-1^, the carbon isotope fractionation between methane and carbon dioxide increases, approaching and eventually overshooting isotopic equilibrium (Figure 3A). This observation is consistent with previous studies^28,62^. In contrast, hydrogen isotope fractionation between methane and water increases slightly with ΔG_r_ and remains far from equilibrium (Figure 3B). Clumped isotopic signatures (Δ^13^CH_3_D and Δ^12^CH_2_D_2_), on the other hand, first approach isotopic equilibrium, but rapidly deviates from equilibrium when ΔG_r_ is around −35 kJ mol^-1^ (Figure 3C and 3D). When the energetics of methanogenesis approaches the thermodynamic limit of microbial metabolism (ΔG_r_ between −35 to −15 kJ mol^-1^), all isotopic signatures drastically change towards equilibrium (Figure 3). This implicates that ΔG_r_ near −35 kJ mol^-1^ is a threshold for isotopic re-equilibration by hydrogenotrophic methanogenesis (Figure 3), above which both bulk and clumped isotopic signatures are overwritten by the equilibrium isotope signatures with increasing ΔG_r_, as suggested previously^26^. Meanwhile, isotopic signatures, in particular ^13C^*ε*_CH4-CO2_, can provide information about the energetics of hydrogenotrophic methanogenesis when ΔG_r_ is more negative than −35 kJ mol^-1^, due to the distinct responses of bulk and clumped isotopic signatures to the variation of ΔG_r_ (Figure 3).

### Natural isotopic tracers for energy availability and methanogenesis pathways

The energetics and metabolic pathways of methanogens can be tracked with the clumped isotope signatures of methane. When paired with environmental context, these enable for a more complete understanding of the controls on methane formation *in situ*. The isotopic signatures of methane are overwritten by re-equilibration process under subsurface conditions, where energy for methanogenesis is very low due to high dissolved methane concentrations^26^. This explains the observation that close-to-equilibrium isotopic signals typically occur under subsurface conditions^19,26,27,47,63–65^. In principle, extremely low-energy availability can result from substrate limitation, yet there is no direct evidence from in-lab cultivation of axenic cultures or cocultures. In contrast to subsurface environments, methanogenesis under surface conditions typically demonstrate more kinetic isotopic signals^20,27,66–68^, indicating a relatively higher Gibbs energy yield of methanogenesis in these environments.

The metabolic pathway of methane formation also determines bulk and clumped isotope signatures. Microbial methanogenesis from methylated organic compounds (methylotrophic, methoxydotrophic, acetoclastic) are characterized by more negative Δ^12^CH_2_D_2_ value than hydrogenotrophic methanogenesis due to the combination of the methyl group and hydrogen from intra-cellular hydrogen carrier (Figure 1)^25,29,36^. We show that hydrogenotrophic methanogenesis at ΔG_r_ around −35 kJ mol^-1^ can generate similar clumped isotopic signatures as acetoclastic, methylotrophic and methoxydotrophic methanogenesis in previous reports (Figure 1)^25,29,59^. Carbon isotope fractionation may provide distinctions. During acetoclastic methanogenesis, the apparent carbon isotope fractionation between methane and CO_2_ (^13*C*^*ε*_CH4-CO2_) ranges from −40 to −20 ‰^25,69,70^, whereas the carbon isotope fractionation is larger for syntrophic methanogenesis with acetate and propionate, ranging from −90 to −80 ‰, as shown in this study (Figure 1A). While distinct *δ*^13^C values can potentially trace hydrogenotrophic methanogenesis operating at low thermodynamic drive, it is influenced by the *δ*^13^C of the carbon source (e.g., CO_2_ or acetate)^48^. This highlights the importance of bulk carbon isotope fractionation paired with clumps when tracing syntrophic methanogenesis at lower energy availabilities.

This study provides experimental constraints for tracing microbial hydrogenotrophic methanogenesis under a range of bioenergetic conditions. We observe a complex response in methane isotopic signatures to the thermodynamic drive available to methanogens. Instead of unidirectionally approaching equilibrium with decreasing Gibbs energy yield, clumped isotope signatures further deviate from equilibrium in an ‘energy niche*’* where the Gibbs energy yield is around −35 kJ mol^-1^. This demonstrates that isotopic re-equilibration by hydrogenotrophic methanogenesis is limited to environments with extremely low energy availability, while the isotopic signatures of microbial methanogenesis stay disequilibrated over a large range of Gibbs energy yield. Additionally, the experimental data provide critical insights into the clumped isotope signatures of methane produced from the syntrophic oxidation of organic acids, highlighting the necessity of both bulk and clumped isotopic analyses in tracing pathways and energetics of microbial methanogenesis.

Microbial methanogenesis is an important terminal step of anaerobic organic matter degradation and a major source of methane in natural and engineered environments^7,71^. The syntrophic degradation of organic compounds, including fatty and aromatic acids, provides essential substrates for methane production and plays a central role in carbon cycling in anoxic ecosystems. These processes are also critical to anaerobic digestion, where the syntrophic oxidation of acetate and propionate supports the conversion of organic wastes into biogas^9,10^. Methane can also be produced directly from substrates such as acetate and methylated compounds through acetoclastic and methylotrophic methanogenesis. The relative contributions of these pathways vary with environmental conditions, substrate availability, and microbial community composition^6,72^. Distinguishing among pathways and constraining the energetics of methanogenesis are therefore essential for understanding methane production in natural environments and optimizing methane management and recovery in engineered systems. Our findings demonstrate that methane bulk and clumped isotope signatures provide valuable constraints on methanogenic pathways and thermodynamics. This study advances a mechanistic understanding of the isotopic fingerprints of methanogenesis and provides a means for deciphering methane cycling across diverse natural and engineered environments.

## ASSOCIATED CONTENT

The following files are available free of charge:

The Supplementary Materials provide additional information on the experimental details, materials, methods, data on microbial growth, isotope data and modeling results. The Supplementary Materials include Figure S1-S3, Table S1-S6 (PDF).

The Python scripts and data frames used in the analysis are on Figshare: https://doi.org/10.6084/m9.figshare.34988050

## AUTHOR INFORMATION

### Author Contributions

Conceptualization: Jiawen Li, William D. Leavitt.

Methodology: Jiawen Li, Anna Schnürer, Maria Westerholm, William D. Leavitt.

Investigation: Jiawen Li, Laetitia Guibourdenche, Dari Bumgarner, Michael Galvin, Xiahong Feng, Annie Kandel, Anna Schnürer, Maria Westerholm.

Visualization: Jiawen Li, Anna Schnürer.

Supervision: Maria Westerholm, Anna Martini, James F. Holden, Edward D. Young, William D. Leavitt.

Funding: Maria Westerholm, James F. Holden, Edward D. Young, William D. Leavitt.

Writing—original draft: Jiawen Li.

Writing—review & editing: Jiawen Li, Laetitia Guibourdenche, Dari Bumgarner, Michael Galvin, Annie Kandel, Anna Schnürer, Maria Westerholm, Anna Martini, James F. Holden, Edward D. Young, William D. Leavitt.

### Funding Sources

Dartmouth College Irving Institute (W.D.L)

Simons Foundation Award 62388 (W.D.L.)

NASA Exobiology Award 80NSSC21K0477 (E.D.Y., W.D.L.)

FORMAS 2024-01865 (A.S.)

### Notes

The authors declare no competing interest.

## ACKNOWLEDGMENT

We thank the team at UC Davis Stable Isotope Facility for carbon isotope measurements on the headspace carbon dioxide and we will deeply miss their absence in future projects. We thank Prof. Cornelia Welte for the help in establishing the collaborations for this work and Dr. Jonathan Gropp for the in-depth discussions on the interpretation of the data and isotopic modeling. Dr. George Cheng is acknowledged for visualization of microbial community structure in the acetate enrichment.

## Notes

### Competing Interest Statement

The authors have declared no competing interest.

