## Supplementary Materials for "Energetic niches of syntrophic methanogens recorded by clumped isotopes"

**Summary:** This document has 12 pages, including 3 figures, 6 tables.

[1] Department of Earth Sciences, Dartmouth College, Hanover, NH, 03755, United States.
[2] Department of Earth, Planetary, and Space Sciences, University of California, Los Angeles, Los Angeles, CA, 90095, United States.
[3] Department of Microbiology, University of Massachusetts Amherst, Amherst, MA, 01003, United States.
[4] Department of Geology & Geophysics, University of Utah, Salt Lake City, UT, 84112, United States.
[5] Department of Molecular Sciences, Swedish University of Agricultural Sciences, Uppsala, SE-750 07, Sweden.
[6] Department of Geology, Amherst College, Amherst, MA, 01002, United States.
[7] Department of Chemistry, Dartmouth College, Hanover, NH, 03755, United States.

Corresponding author:

*Jiawen Li* – Department of Earth Sciences, Dartmouth College, NH, 03755, United States.

Present Address

† Department of Ocean Sciences, Rosenstiel School of Marine, Atmospheric, and Earth Science, University of Miami, Miami, FL 33149, United States.

### Contents


1. **Supplementary Figures**

   **Figure S1.** Sequencing results of the bioreactors at SLU.

   **Figure S2.** Headspace gas amounts in monoculture and coculture experiments at Dartmouth.

   **Figure S3.** Isotopic model results with MCR I.

2. **Supplementary Tables**

   **Table S1.** Growth conditions of the bioreactors at SLU.

   **Table S2.** Headspace gas amounts and calculated Gibbs free energy yield (ΔG) of methanogenesis.

   **Table S3.** Summary of methane isotope measurements.

   **Table S4.** Substrate isotope signatures.

   **Table S5.** Methane production rates in the experiments.

   **Table S6.** Hydrogen and carbon isotope fractionation factors of methanogenesis experiments.

3. **Supplementary Texts**

   **Text S1.** C-P medium recipe.

   **Text S2.** DSM 282 medium recipe.

   **Text S3.** Estimation of substrate consumption based on stoichiometry

#### Supplementary Figures

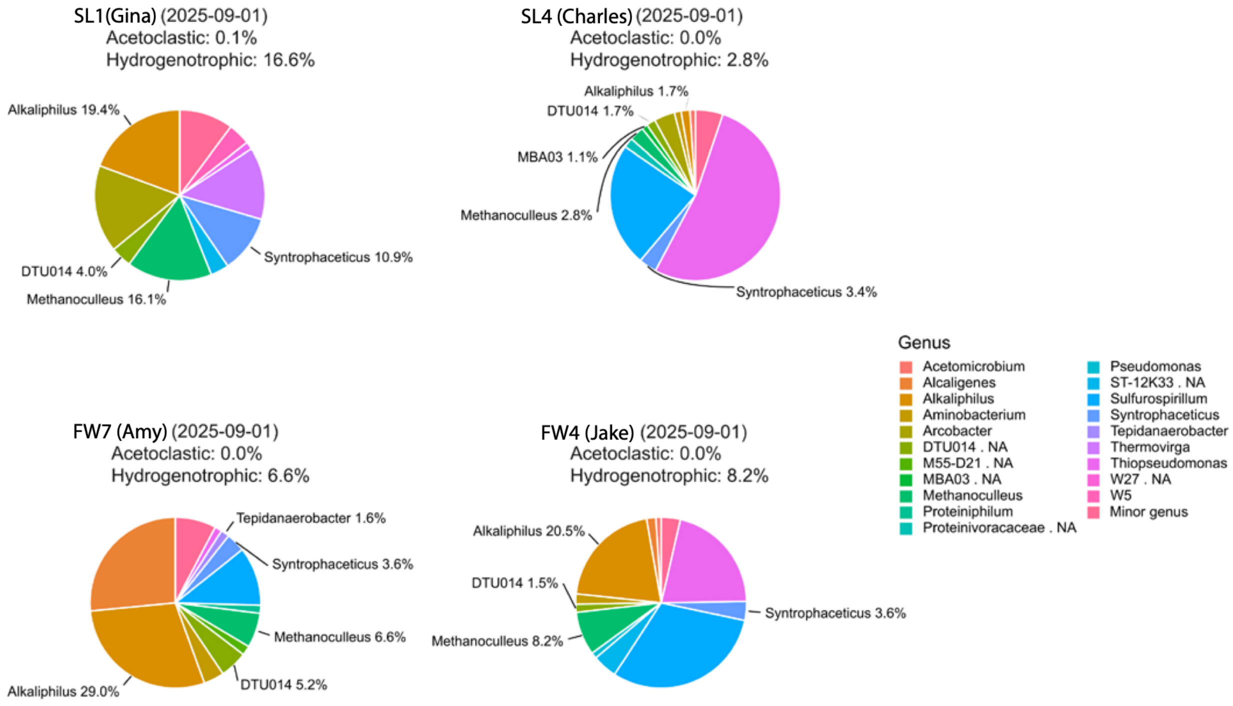


**Figure S1. Sequencing results of the bioreactors at SLU.**

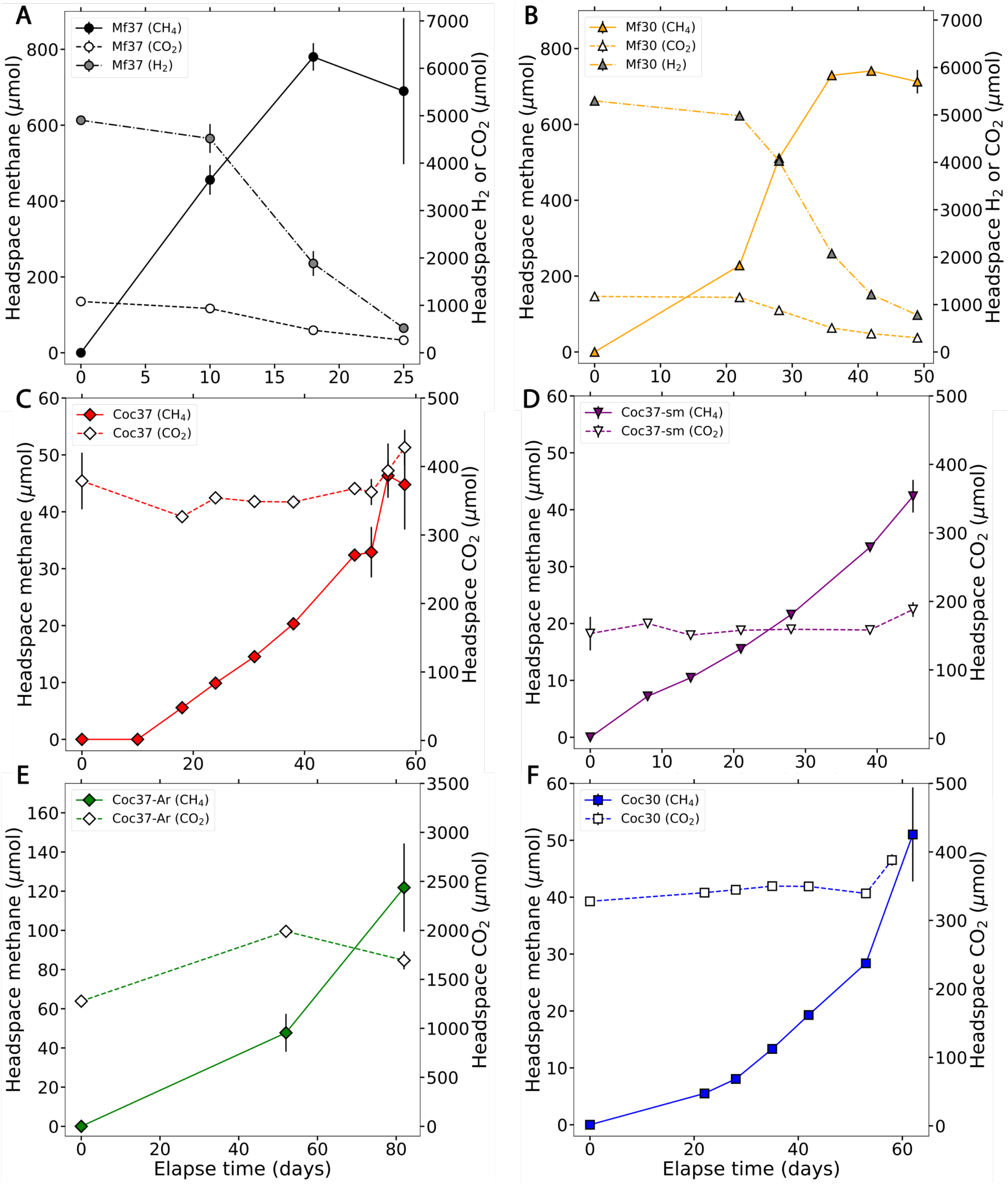


**Figure S2.** Headspace gas amounts in monoculture and coculture experiments at Dartmouth. Each panel represents one experiment: (A) Mf37; (B) Mf30; (C) Coc37; (D) Coc37-sm; (E) Coc37-Ar; (F) Coc30. For monoculture methanogenesis with $H_2/CO_2$ (panels A-B), hydrogen gas, carbon dioxide and methane amounts were measured during the experiment. For coculture methanogenesis (panels C-F), Hydrogen gas amount is below the detection limit of routine measurements. Therefore, only carbon dioxide and methane amounts were routinely measured, and the trace amount of hydrogen gas was quantified at the time of sacrifice.

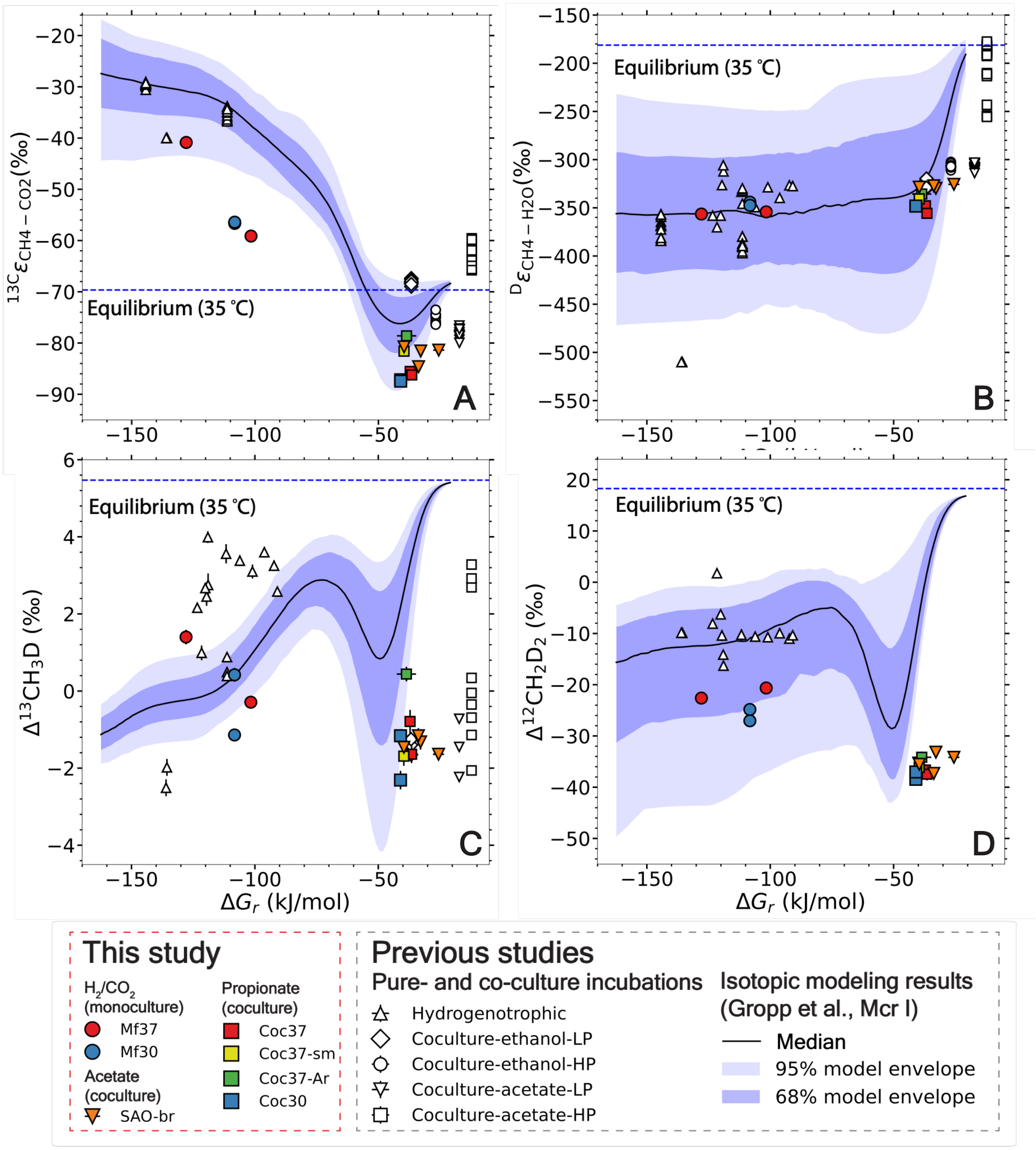


**Figure S3.** Variations of isotope fractionation and isotopic signatures with the $\Delta G_r$ of hydrogenotrophic methanogenesis: (A) $^{13C}\varepsilon_{CH4\text{-}CO2}$; (B) $^{D}\varepsilon_{CH4\text{-}H2O}$; (C) $\Delta^{13}CH_3D$ and (D) $\Delta^{12}CH_2D_2$. Similar to Figure 3, the median of 200 model simulations with MCR I are shown by the black lines, and the deep and light blue denotes the envelopes of 68 % and 95 % of the results.

#### Supplementary Tables

**Table S1. Growth conditions of the bioreactors in SLU.**

| Bioreactors | pH | TAN / g $L^{-1}$ | FAN / mg $L^{-1}$ |
|---|---|---|---|
| SL1 (Gina) | 8.1 | 1 | 139 |
| FW4 (Jake) | 7.9 | 4 | 370 |
| SL4 (Charles) | 7.9 | 4 | 370 |
| FW7 (Amy) | 7.8 | 7 | 524 |

**Table S2. Headspace gas amounts and calculated Gibbs free energy yield ($\Delta G_r$) of hydrogenotrophic methanogenesis.** All uncertainties represent 1-σ errors.

| Experiments | Incubation time (days) | $H_2$ (μmol) | $CO_2$ (μmol) | $CH_4$ (μmol) | Vol (mL) | Temp (°C) | $\Delta G_r$ (kJ/mol) |
|---|---|---|---|---|---|---|---|
| **Dartmouth*** | | | | | | | |
| Mf37_T1 | 10 | 4519.7 ± 305.8 | 934.2 ± 94.7 | 455.9 ± 39.2 | 110 | 37 | -127.9 ± 0.8 |
| Mf37_T2 | 18 | 1882.7 ± 261.6 | 475.1 ± 44.7 | 780.1 ± 36.3 | 110 | 37 | -115.8 ± 1.5 |
| Mf37_T3 | 25 | 529.7 ± 38.0 | 283.0 ± 19.8 | 689.9 ± 27.1 | 110 | 37 | -101.7 ± 0.8 |
| Mf30 | 49 | 778.3 ± 99.4 | 299.5 ± 33.8 | 712.6 ± 31.0 | 110 | 30 | -108.3 ± 1.3 |
| Coc37_T1 | 52 | 0.54 ± 0.06 | 362.7 ± 19.0 | 32.9 ± 4.4 | 108.5 | 37 | -39.2 ± 1.2 |
| Coc37_T2 | 55 | 0.47 ± 0.06 | 394.0 ± 39.7 | 46.4 ± 2.8 | 108.5 | 37 | -37.1 ± 1.4 |
| Coc37_T3 | 58 | 0.43 ± 0.06 | 427.8 ± 25.9 | 44.8 ± 7.9 | 108.5 | 37 | -36.5 ± 1.5 |
| Coc37-sm | 45 | 0.13 ± 0.01 | 188.2 ± 10.8 | 51.0 ± 8.3 | 18.5 | 37 | -39.6 ± 0.9 |
| Coc37-Ar | 85 | 0.48 ± 0.18 | 1694.0 ± 91.7 | 121.8 ± 22.5 | 108.5 | 37 | -38.6 ± 3.9 |
| Coc30 | 62 | 0.45 ± 0.06 | 388.1 ± 8.3 | 42.4 ± 2.9 | 108.5 | 30 | -41.0 ± 1.4 |
| **SLU** | | | | | | | |
| SAO-br-SL1 | n.a. | 0.48 ± 0.04 | 1004.6 ± 149.6 | 1808.1 ± 126.0 | 118 | 37 | -33.7 ± 0.9 |
| SAO-br-SL4 | n.a. | 0.18 ± 0.04 | 1442.7 ± 124.5 | 1304.1 ± 61.8 | 118 | 37 | -25.6 ± 2.3 |
| SAO-br-FW4 | n.a. | 0.41 ± 0.04 | 1081.1 ± 211.7 | 1460.0 ± 201.6 | 118 | 37 | -32.9 ± 1.2 |
| SAO-br-FW7 | n.a. | 0.72 ± 0.04 | 1279.2 ± 63.2 | 1129.3 ± 123.7 | 118 | 37 | -39.6 ± 0.6 |
| SAO-batch-SL1 | 42 | n.d.† | 438.59 ± 60.31 | 508.68 ± 36.59 | 78 | 37 | n.a. ‡ |
| SAO-batch-SL4 | 42 | n.d. | 240.30 ± 13.65 | 142.83 ± 7.24 | 78 | 37 | n.a. |
| SAO-batch-FW4 | 42 | n.d. | 329.04 ± 5.82 | 271.47 ± 7.97 | 78 | 37 | n.a. |
| SAO-batch-FW7 | 42 | n.d. | 218.54 ± 12.93 | 252.49 ± 22.54 | 78 | 37 | n.a. |

* There were three time points of sacrifice in Mf37 and Coc37 experiments, which are noted as T1, T2 and T3.

† n.d. indicates that the hydrogen gas is below detection limit.

‡ n.a. indicates the estimation of Gibbs free energy yield is not available due to below-detection-limit hydrogen gas amount.

**Table S3. Summary of methane isotope measurements.** All uncertainties represent 1-σ errors.

| Samples | Incubation time (days) | $\delta^{13}C$ ($CH_4$) (‰) | δD ($CH_4$) (‰) | $\Delta^{13}CH_3D$ (‰) | $\Delta^{12}CH_2D_2$ (‰) |
|---|---|---|---|---|---|
| **Dartmouth** | | | | | |
| Mf37-T1-3 | 10 | -56.395 ± 0.004 | -390.466 ± 0.036 | 1.40 ± 0.19 | -22.61 ± 0.91 |
| Mf37-T3-2 | 25 | -58.420 ± 0.007 | -388.503 ± 0.027 | -0.29 ± 0.14 | -20.64 ± 0.90 |
| Mf30-1 | 49 | -58.661 ± 0.040 | -379.257 ± 0.026 | -1.14 ± 0.14 | -27.04 ± 0.80 |
| Mf30-2 | 49 | -58.410 ± 0.004 | -382.415 ± 0.033 | 0.42 ± 0.18 | -24.84 ± 0.98 |
| Coc37-T2-2 | 55 | -92.529 ± 0.005 | -382.405 ± 0.038 | -0.79 ± 0.30 | -36.65 ± 1.10 |
| Coc37-T3-2 | 58 | -93.076 ± 0.027 | -389.338 ± 0.040 | -1.64 ± 0.23 | -37.46 ± 1.21 |
| Coc37-sm-1 | 45 | -88.715 ± 0.028 | -375.784 ± 0.033 | -1.69 ± 0.25 | -36.14 ± 1.08 |
| Coc37-Ar-1 | 85 | -91.007 ± 0.022 | -405.168 ± 0.041 | 0.44 ± 0.19 | -34.16 ± 1.09 |
| Coc30-2 | 62 | -94.066 ± 0.007 | -382.575 ± 0.024 | -1.16 ± 0.17 | -38.46 ± 0.89 |
| Coc30-3 | 62 | -94.332 ± 0.014 | -382.624 ± 0.048 | -2.31 ± 0.24 | -37.03 ± 1.32 |
| **UMass** | | | | | |
| Mj-fm-1 | 0.4 | -79.282 ± 0.005 | -391.201 ± 0.026 | 1.62 ± 0.14 | -24.65 ± 0.88 |
| Mj-fm-3 | 0.5 | -74.011 ± 0.005 | -372.723 ± 0.024 | 2.61 ± 0.24 | -25.82 ± 0.85 |
| **SLU** | | | | | |
| SLU-br-SL4 | n.a. | -85.878 ± 0.004 | -373.813 ± 0.027 | -1.64 ± 0.14 | -34.17 ± 0.75 |
| SLU-br-FW7 | n.a. | -85.528 ± 0.005 | -375.301 ± 0.034 | -1.46 ± 0.18 | -35.53 ± 0.88 |
| SLU-br-FW4 | n.a. | -85.656 ± 0.004 | -377.920 ± 0.039 | -1.32 ± 0.20 | -33.17 ± 0.84 |
| SLU-br-SL1 | n.a. | -87.703 ± 0.004 | -375.035 ± 0.038 | -1.16 ± 0.20 | -37.31 ± 0.87 |
| SAO-batch-SL1 | 42 | -92.384 ± 0.006 | -377.612 ± 0.019 | -2.02 ± 0.33 | -37.58 ± 0.94 |
| SAO-batch-SL4 | 42 | -89.934 ± 0.005 | -378.997 ± 0.035 | -1.44 ± 0.21 | -30.56 ± 1.42 |
| SAO-batch-FW4 | 42 | -92.538 ± 0.006 | -382.8 ± 0.032 | -1.78 ± 0.27 | -38.13 ± 1.26 |
| SAO-batch-FW7 | 42 | -88.697 ± 0.006 | -371.024 ± 0.023 | -2.32 ± 0.37 | -36.92 ± 1.03 |

**Table S4. Substrate isotope signatures.**

| Samples | $\delta D_{H2O}$ (‰) | $\delta^{13}C_{CO2}$ (‰) |
|---|---|---|
| **Dartmouth** | | |
| Mf37-T1-3 | -52.85 ± 0.15 | -16.20 ± 0.07 |
| Mf37-T3-2 | -53.07 ± 0.06 | 0.75 ± 0.07 |
| Mf30-1 | -53.58 ± 0.08 | -2.12 ± 0.07 |
| Mf30-2 | -53.58 ± 0.08 | -2.12 ± 0.07 |
| Coc37-T2-2 | -52.88 ± 0.07 | -7.63 ± 0.07 |
| Coc37-T3-2 | -52.19 ± 0.08 | -7.49 ± 0.07 |
| Coc37-sm-1 | -53.15 ± 0.12 | -7.76 ± 0.07 |
| Coc37-Ar-1* | -104.6 ± 0.20 | -13.47 ± 0.07 |
| Coc30-2 | -52.81 ± 0.13 | -7.55 ± 0.07 |
| Coc30-3 | -52.81 ± 0.13 | -7.55 ± 0.07 |
| **UMass** | | |
| Mj-fm-1 | -50.70 ± 0.20 | -7.75 ± 0.07 |
| Mj-fm-3 | -50.70 ± 0.20 | -7.75 ± 0.07 |
| **SLU** | | |
| SLU-br-SL4 | -71.40 ± 0.20 | -4.88 ± 0.07 |
| SLU-br-FW7 | -69.60 ± 0.20 | -5.19 ± 0.07 |
| SLU-br-FW4 | -71.90 ± 0.20 | -4.46 ± 0.07 |
| SLU-br-SL1 | -71.20 ± 0.20 | -3.36 ± 0.07 |

* This experiment was conducted at the University of Utah and therefore has different hydrogen and carbon isotope values for water and $CO_2$ than the other experiments at Dartmouth College.

**Table S5. Methane production rates across different experiments.** The production rates of the batch culturing of SAO enrichment at SLU is not tracked over the course of the experiments and thus is not shown on this table.

| Experiment | Days | Total Methane (µmol) | Rate* (µmol $mL^{-1}$ $day^{-1}$) | Culture Volume (mL) |
|---|---|---|---|---|
| **Dartmouth** | | | | |
| Mf37 | 18 | 780.06 ± 36.28 | 0.8667 ± 0.0403 | 50 |
| Mf30 | 42 | 712.63 ± 30.98 | 0.3393 ± 0.0148 | 50 |
| Coc37 | 55 | 46.36 ± 2.76 | 0.0164 ± 0.0010 | 51.5 |
| Coc37-sm | 45 | 42.37 ± 2.87 | 0.0183 ± 0.0012 | 51.5 |
| Coc37-Ar | 82 | 121.83 ± 22.52 | 0.0288 ± 0.0053 | 51.5 |
| Coc30 | 62 | 51.02 ± 8.27 | 0.0160 ± 0.0026 | 51.5 |
| **UMass** | | | | |
| Mj-fm-1 | 0.375† | 69.09 ± 9.84 | 7.369 ± 1.049 | 25 |
| Mj-fm-3 | 0.528 | 109.57 ± 15.98 | 8.302 ± 1.211 | 25 |

* Rate represents the volumetric methane production rate over the course of the designated days at the exponential growth phase. Uncertainties are presented as the 1 standard deviation.
† Incubations with formate were completed within 1 day, but the rates are shown in the unit of µmol $mL^{-1}$ $day^{-1}$ for consistency.

**Table S6. Hydrogen and carbon isotope fractionation factors of methanogenesis experiments.** All uncertainties are 1-σ errors.

| Samples | ${}^{D}\alpha_{CH4\text{-}H2O}$ | ${}^{13C}\alpha_{CH4\text{-}CO2}$ | ${}^{D}\varepsilon_{CH4\text{-}H2O}$ (‰) | ${}^{13C}\varepsilon_{CH4\text{-}CO2}$ (‰) |
|---|---|---|---|---|
| **Dartmouth** | | | | |
| Mf37-T1 | 0.6435 ± 0.00011 | 0.9591 ± 0.00007 | -356.45 ± 0.11 | -40.86 ± 0.07 |
| Mf37-T3 | 0.6458 ± 0.00005 | 0.9409 ± 0.00007 | -354.23 ± 0.05 | -59.13 ± 0.07 |
| Mf30-1 | 0.6559 ± 0.00006 | 0.9433 ± 0.00008 | -344.12 ± 0.06 | -56.66 ± 0.08 |
| Mf30-2 | 0.6525 ± 0.00007 | 0.9436 ± 0.00007 | -347.45 ± 0.07 | -56.41 ± 0.07 |
| Coc37-T2 | 0.6521 ± 0.00006 | 0.9144 ± 0.00007 | -347.93 ± 0.06 | -85.55 ± 0.06 |
| Coc37-T3 | 0.6443 ± 0.00007 | 0.9138 ± 0.00007 | -355.71 ± 0.07 | -86.23 ± 0.07 |
| Coc37-sm | 0.6593 ± 0.00009 | 0.9184 ± 0.00007 | -340.74 ± 0.09 | -81.59 ± 0.07 |
| Coc37-Ar | 0.6643 ± 0.00016 | 0.9214 ± 0.00007 | -335.68 ± 0.16 | -78.60 ± 0.07 |
| Coc30-1 | 0.6518 ± 0.00010 | 0.9126 ± 0.00007 | -348.20 ± 0.10 | -87.44 ± 0.07 |
| Coc30-2 | 0.6519 ± 0.00009 | 0.9128 ± 0.00007 | -348.15 ± 0.09 | -87.17 ± 0.06 |
| **UMass** | | | | |
| Mj-fm-1 | 0.6413 ± 0.00014 | 0.9279 ± 0.00007 | -358.69 ± 0.14 | -72.09 ± 0.07 |
| Mj-fm-2 | 0.6608 ± 0.00014 | 0.9332 ± 0.00007 | -339.22 ± 0.14 | -66.78 ± 0.07 |
| **SLU** | | | | |
| SLU-br-SL4 | 0.6743 ± 0.00015 | 0.9186 ± 0.00007 | -325.67 ± 0.15 | -81.39 ± 0.06 |
| SLU-br-FW7 | 0.6714 ± 0.00015 | 0.9192 ± 0.00007 | -328.57 ± 0.15 | -80.76 ± 0.06 |
| SLU-br-FW4 | 0.6703 ± 0.00015 | 0.9184 ± 0.00007 | -329.73 ± 0.15 | -81.56 ± 0.06 |
| SLU-br-SL1 | 0.6729 ± 0.00015 | 0.9154 ± 0.00006 | -327.13 ± 0.15 | -84.63 ± 0.06 |

**Supplementary Texts**

**Text S1. C-P medium recipe**

The C-P medium is a bicarbonate-buffered basal solution containing the following component (per L): 0.53 g $Na_2HPO_4 \cdot 2H_2O$; 0.41 g $KH_2PO_4$; 0.3 g $NH_4Cl$; 0.3 g NaCl; 0.11 g $CaCl_2 \cdot 2H_2O$; 0.1 g $MgCl_2 \cdot 6H_2O$; 0.5 mg resazurin; 7.5 μM $FeCl_2$; 1 μM $H_3BO_4$; 0.5 μM $ZnCl_2$; 0.1 μM $CuCl_2$; 0.5 μM $MnCl_2$; 0.5 μM $CoCl_2$; 0.1 μM $NiCl_2$; 0.1 μM $Na_2SeO_3$; 0.1 μM $Na_2WO_4$; and 0.1 μM $Na_2MoO_4$. The medium was autoclaved and supplemented with (per L): 4 g $NaHCO_3$, 0.48 g $Na_2S \cdot 9H_2O$, 0.5 g cysteine and 1 mL of 1000× Wolin's vitamin solution post-autoclave. All the post-autoclave supplements were made anoxic and sterilized by autoclaving except for the vitamin solution, which was filter sterilized. Specifically, the 1000× vitamin solution contains (per L): Biotin 20 mg; Folic acid 20 mg; Pyridoxine hydrochloride 100 mg; Thiamine HCl 50 mg; Riboflavin 50 mg; Nicotinic acid 50 mg; Calcium D-(+)-pantothenate 50 mg; Vitamin $B_{12}$ 1 mg; p-Aminobenzoic acid 50 mg and (DL)-alpha-Lipoic acid 50 mg. For the cultivation of *S. fumaroxidans*, the medium was supplemented with 20 mM sodium fumarate, while for the cultivation of the co-culture of *S. fumaroxidans* and *M. formicicum*, the medium was supplemented with 30 mM of propionate. The final pH of the medium was adjusted to 7.0-7.2 with 1 M NaOH or 1 M HCl.

**Text S2. DSM 282 medium recipe**

The DSM 282 growth medium contained the following (per L): 780 mL of $ddH_2O$, 200 mL of 5× salts (containing per L: 150 g NaCl, 20.5 g $MgCl_2 \cdot 6H_2O$, 17 g $MgSO_4 \cdot 7H_2O$, 1.65 g KCl, 1.25 g $NH_4Cl$, 0.7 g $CaCl_2 \cdot 2H_2O$, 0.7g $K_2HPO_4$), 1 g $NaHCO_3$, 1 g sodium formate, 10 mL DSM 141 trace vitamins (containing per L: 2 mg biotin, 2 mg folic acid, 10 mg pyridoxine-HCl, 5 mg thiamine-HCl·$2H_2O$, 5 mg riboflavin, 5 mg nicotinic acid, 5 mg D-pantothenate, 0.1 mg vitamin $B_{12}$, 5 mg p-aminobenzoic acid, 5 mg lipoic acid (6,8-thioctic acid)), 10 mL 141 trace minerals (containing per L: 1.5 g nitrilotriacetic acid, 3 g $MgSO_4 \cdot 7H_2O$, 0.5 g $MnSO_4 \cdot H_2O$, 1 g NaCl, 1 g $FeSO_4 \cdot 7H_2O$, 0.18 g $CoSO_4 \cdot 7H_2O$, 0.1 g $CaCl_2 \cdot 2H_2O$, 0.18 $ZnSO_4 \cdot 7H_2O$, 0.01 g $CuSO_4 \cdot 5H_2O$, 0.02 $KAl(SO_4)_2 \cdot 12H_2O$, 0.01 g $H_3BO_3$, 0.01 g $Na_2MoO_4 \cdot 2H_2O$, 0.025 g $NiCl_2 \cdot 6H_2O$), 1 g resazurin, 0.1 mL 0.1 % $Na_2WO_4 \cdot 2H_2O$/0.1 % $Na_2SeO_4$. The medium was pH balanced to 6.00 ± 0.05.

**Text S3. Estimation of substrate consumption**

The average initial headspace $H_2$ amounts were 4904 and 5296 μmol for the cultivation of monoculture Mf under 37 and 30 ºC, respectively. At the last time point of sampling, the headspace $H_2$ amounts dropped to 520 and 778 μmol under 37 and 30 ºC, respectively. These corresponded to 89 and 85 % of the initial $H_2$ under these two growth temperatures.

The consumption of propionate during the cultivation of coculture Mf+Sf was estimated based on the stoichiometry of the syntrophic methanogenesis on propionate (Table 1). The maximum amount of methane production is 1136 μmol, assuming a full consumption of the 30 mM propionate supplemented. The actual methane produced by the Mf+Sf coculture, however, ranged from 32.9 to 121.8 μmol (Table S2). These corresponded to only 3-10 % of the initial propionate supplemented.
